# Scanning transcriptomes for nonlinear, domain-level similarities using hmSEEKR

**DOI:** 10.64898/2026.07.03.736302

**Authors:** Shuang Li, Daniel A. Sprague, Quinn E. Eberhard, Samuel P. Boyson, Alain Laederach, J. Mauro Calabrese

## Abstract

Long noncoding RNAs (lncRNAs) play roles in gene regulation across kingdoms of life. However, lncRNAs with related functions often lack linear sequence similarity, making it difficult to leverage studies of one lncRNA to inform the understanding of others. We describe a *k*-mer-based hidden Markov model, hmSEEKR, that enables the scanning of transcriptomes for regions of non-linear sequence similarity to a query domain, without prior knowledge of where within the transcriptome the similarities may be located. When individual lncRNA domains were used as search features, hmSEEKR successfully identified regions in other RNAs that harbor non-linear sequence similarity and bind similar sets of proteins. Applying hmSEEKR to transcriptome-wide searches, we found that certain domains within the lncRNAs *XIST*, *NEAT1*, and *MALAT1* exhibited widespread regional similarity to both lncRNA and protein-coding genes, while others were more unique, exhibiting similarity to ∼100 genes or fewer. Combinatorial searches uncovered RNAs containing sequential matches to core functional domains of *XIST* and *NEAT1*, and eCLIP-inferred protein-interaction networks within these RNAs more closely resembled those of *XIST* and *NEAT1*, respectively, than would be expected by chance, suggesting the searches recovered RNAs with similar biological properties. Finally, within annotated sets of cis-activating and cis-repressive lncRNAs, we observed opposing enrichments for similarity to domains associated with transcription-promoting complexes and heterogeneous nuclear ribonucleoprotein (hnRNP) binding, respectively, suggesting the enriched sequences may contribute to regulatory functions. hmSEEKR can be applied with minimal training data and enables the a priori discovery of RNA domains that share nonlinear similarity, offering a sequence-informed approach to discover functional elements within noncoding transcriptomes.

## Introduction

Long noncoding RNAs (lncRNAs) play important roles in the regulation of many biological processes. However, the majority of annotated lncRNAs lack ascribed functions. For example, current annotations suggest that the human genome can produce over 190000 different lncRNA transcripts from ∼36000 loci (1). Yet, to date, a much smaller number, on the order of hundreds, have been linked to a specific molecular or cellular function (2–4).

Ascribing function to newly identified lncRNAs remains challenging. Unlike protein-coding genes, lncRNAs are not constrained by codon usage, evolve rapidly, and achieve function by employing structures and proteins in ways that are not well-understood. Thus, lncRNAs with related functions rarely harbor significant stretches of linear sequence similarity, rendering ineffective a large set of sequence comparison tools that help ascribe function to protein-coding genes, and compounding the difficulty of inferring function of one lncRNA from the study of another (5, 6).

We previously developed a simple non-linear comparison framework, called SEquence Evaluation through *K*-mer Representation (SEEKR), which enables sequences to be compared by the relative abundance of the *k*-mers contained within them. Using SEEKR, we demonstrated that lncRNAs with related functions can harbor related *k*-mer contents and that *k*-mer content can be used to predict protein-binding to lncRNAs and their cellular localization (5–8). These studies and others suggest nonlinear comparison frameworks such as SEEKR can provide meaningful information about similarities between lncRNAs in cases where linear alignments may fail.

However, a shortfall of the SEEKR algorithm is that without user input, SEEKR is unable to consider positional information in similarity evaluations. Many lncRNAs exert functions through local domains that have distinct *k*-mer contents that can become averaged away in whole-transcript similarity searches (2–4). Thus, when using SEEKR to identify regional similarities to functional domains of interest, the set of target lncRNAs being searched must be first broken up into predefined fragments by the end-user. Yet in a set of functionally uncharacterized target lncRNAs, it would be unclear where the boundaries of putative functional domains might be located, increasing the risk that fragmentation may split domains and cause regional similarity to be missed (7, 8).

The Hidden Markov Model (HMM) is a statistical framework that has been broadly employed in the biological sciences to identify hidden patterns from observable sequences (9–11). It offers a way to identify regional similarity to a lncRNA domain of interest without predefined knowledge of where similar domains may be located in other lncRNA(s). Herein, we describe the development and implementation of a generalizable two-state HMM, hmSEEKR, which can detect regions of RNA whose *k*-mer contents are more similar to a “query” (e.g. a domain of interest) than a “null” set of background sequences (e.g. the transcriptome). We demonstrate that hmSEEKR can identify domains in target RNAs that bind similar sets of proteins as their corresponding queries. We apply hmSEEKR to identify domains throughout the transcriptome that exhibit significant similarity to regions across three conserved lncRNAs, *XIST*, *NEAT1*, and *MALAT1*, and observe prevalent similarity between the three lncRNAs and both protein-coding and lncRNA genes. We identify RNAs that harbor sequential, domain-based similarity to minimal functional versions of *XIST* and *NEAT1*, and show that these RNAs exhibit evidence of interacting with proteins in similar patterns, supporting the possibility that hmSEEKR identified RNAs that have analogous functions. Finally, we identify that RNA domains with characterized molecular functions exhibit preferential enrichment in lncRNAs proposed to locally activate and repress transcription, establishing footholds for future experimental studies. A strength of hmSEEKR is that the algorithm can be trained on standard computing resources using minimal data – even down to a single domain in a single lncRNA – then rapidly employed to identify regions in other RNAs that harbor similar *k*-mer contents.

## Results

### Overview of hmSEEKR

#### Scanning a pool of sequences for similarity to a query domain

Accumulating evidence suggests that different lncRNAs can carry out similar functions using shared mechanisms, particularly in the context of gene regulation (2, 12–14). Thus, functional domains identified in one lncRNA could serve as exemplars to discover domains with analogous functions in others (Figure 1A). Under this premise, we implemented “hmSEEKR”, a two-state HMM designed to detect regions of RNA whose *k*-mer content is more similar to a query “Q” (a domain of interest) than a null model “N” (the *k*-mer content of a background set of sequences such as the transcriptome; Figure 1B). In the hmSEEKR approach, *k*-mer frequencies for Q and N are calculated by tiling across each query and null sequence at a user-specified *k*-mer length, one nucleotide at a time, then dividing resultant *k*-mer counts by the total length of the sequences in the query and null sets, respectively (Figure 1B, STEP 1). *k*-mer frequencies are then used to populate emission matrices for the Q and N states of the HMM. After the end-user decides upon probabilities of transitioning between Q and N (following guidance described in the subsequent section; Figure 1B, STEP 2), each sequence in a user-defined search pool is scanned using *k* as the window size and a single nucleotide as the step size, and states are defined via the Viterbi algorithm (Figure 1B, STEP 3; (9, 10)).

**Figure 1.**
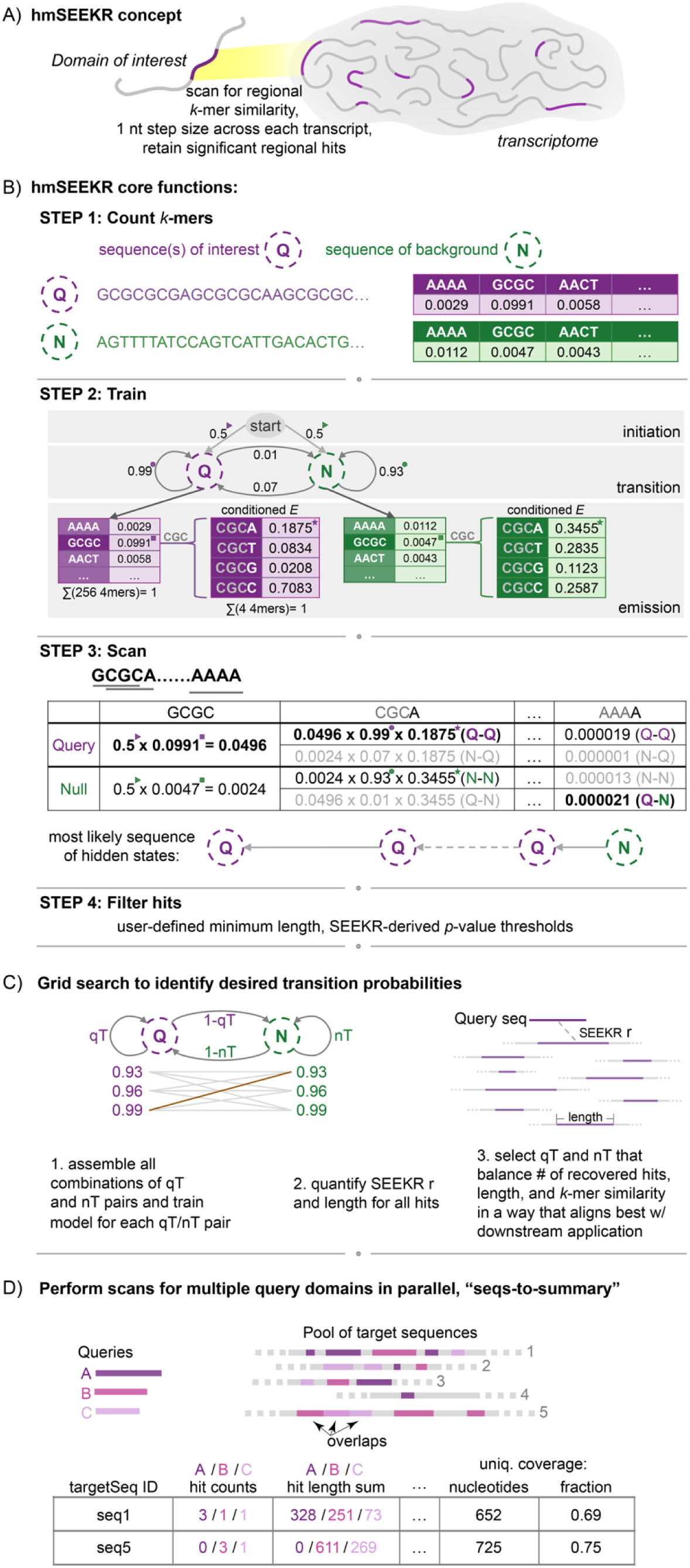
Overview of hmSEEKR. **(A)** Overview of hmSEEKR concept. The basic principles of *k*-mer counting using SEEKR were applied to a hidden Markov model, to enable the scanning of transcriptomes for domain-level *k*-mer similarities (5–8). **(B)** Overview of hmSEEKR algorithm. A single domain or a group of related sequences of interest serves as the “Query”, and a larger set of sequences serve as the “Null”. With user-defined transition parameters qT and nT, the algorithm scans a set of user-defined target sequences, one *k*-mer at a time using a single nucleotide step size and identifies regions of consecutive *k*-mers that are more likely to be classified as Query versus Null (i.e. “hits”). Hits can then be filtered by length and SEEKR-derived *p* values. In the example calculation shown at the bottom of (A): filled triangles mark the initiation probabilities of the Q or N states, which are set by default to 0.5; filled squares mark initial *k*-mer emission probability; filled circles mark qT/nT values; asterisks mark conditioned emission probability. **(C)** Grid search to empirically determine suitable qT and nT values for a given query and set of background sequences. **(D)** “seqs-to-summary” search to enable parallel scans of multiple query domains against a shared set of target sequences. See also Table S1.

*k*-mer counting using a single-nucleotide step-size has proven to be a robust strategy for detecting similarities that underlie shared protein-binding events between lncRNAs (5–8, 13). However, *k*-mer frequencies obtained via step-sizes less than the specified *k*-mer length *k* are not completely independent, making them at odds with the data typically input into an HMM (9, 10). To better align the strengths of *k*-mer counting by single-nucleotide steps with the independent counting principles of an HMM, we implemented a “conditioned emission” approach to determine the probability of each *k*-mer arising from the Q state and N states during hmSEEKR scans (Figure 1B, STEP 3). Under the conditioned approach, emission matrices from STEP1 are reduced to include only the four *k*-mers that can occur at the given position based on the prior *k*-mer in the sequence being searched. For example, using a single nucleotide step, if the prior *k*-mer were ‘GCGC’, the subsequent *k*-mer must begin with ‘CGC’ (Figure 1B, STEP 3). Thus, “conditioned” emission matrices for the Q and N states are calculated as the frequency of each *k*-mer in the original emission matrices divided by the sum of the frequencies of all four possible *k*-mers in the original emission matrices (Figure 1B, STEP 3).

Lastly, following the standard hidden Markov model approach, cumulative Q and N probabilities are calculated for each *k*-mer forwardly through the length of each searched sequence. The most likely state for each *k*-mer along the length of each sequence (either Q or N) is determined by assigning the state of the last *k*-mer with the highest probability and tracing backward. Regions of consecutive *k*-mers assigned to the Q state are retained as “hits”, which can be filtered by parameters set by the end-user, including hit length and SEEKR-derived *p* values that estimate the likelihood of observing the same level of *k*-mer similarity between the hit and the query by chance (Figure 1B; (7)).

#### Defining transition parameters for hmSEEKR scans

Probabilities for transitioning between states must be defined prior to any HMM search. For this purpose, the Baum-Welch algorithm is often used, which employs expectation-maximization to find transition parameters that best recover a set of observed features from a training dataset (9, 10). However, where lncRNAs are concerned, suitable training datasets to optimize transition parameters between Q and N states may not exist. This is because in a single-domain search, transition probabilities derived by optimally recovering the query domain from the host lncRNA are not guaranteed to be generalizable throughout the pool of search sequences, and may not even be representative. For example, the expanded Repeat D region spans about one-third of the length of the human *XIST* transcript (15). While Repeat D-like regions may exist in other lncRNAs, it would be presumptuous to expect those regions to be of proportional length and position within in their host lncRNAs (8, 13).

An alternative to Baum-Welch is to derive transition probabilities empirically, by searching a “grid” of transition parameters to scan a set of known sequences and evaluate the output to identify which parameters perform best on a reference task. In the case of hmSEEKR, a reference task generalizable to any search is to identify the transition probabilities that maximize the SEEKR-derived Pearson’s *r* values between the query domain and the set of hits from a training dataset; these *r* values provide a direct measure of the non-linear sequence similarity that in principle, the end-user would be trying to maximally recover over an hmSEEKR hit (6, 7). To this end, we implemented a grid-search function to provide a semi-automated framework to empirically derive transition parameters for any sequence search (Figure 1C). Over a grid of parameters defined by the end-user, the grid-search function scans a search pool for HMM hits to a query domain(s) and then outputs summary information associated with each condition tested. The summary information includes the number of hits identified, and the median, average, and standard deviation of the both the hit lengths and the SEEKR-derived Pearson’s r values associated with hits. For a given grid search, a sequence pool would ideally be large enough to comprise a diversity of sequences and yield on the order of hundreds of hits to the query domain. In our work below, we optimized all searches against the set of de-duplicated human lncRNAs >500 nucleotides (nts) in length from the GENCODE v47 canonical collection (16).

In developing the grid search function, we found it useful to set a minimum tolerated length for hits and evaluate transition parameters by looking at statistics of the top 50 hits ranked by SEEKR-derived r value to the query domain(s). In our domain searches below, our preferred transition probabilities were often those that maximized the median SEEKR r value among the top 50 hits and yielded a total count of length-filtered hits that was greater than the smaller of 10000 or the median number of length-filtered hits among all tested combinations of transition probabilities. This heuristic approach aimed to balance selectivity with discovery power. Once selected, the set of transition parameters for a given value of *k* can then be employed by the end-user to search for domains that are similar to the query of interest.

Depending on the number of parameters being searched, the size of the training dataset, and the number of query domains being studied, we observed that grid searches can be time consuming and computationally expensive. Over the course of our work below, we optimized parameters for 78 separate query domains and observed that based on the heuristic described above, our two most-commonly selected transition parameters at *k*-mer lengths *k* = 4, 5, and 6 were qT=0.99 and nT=0.93, qT=0.99 and nT=0.90, and qT=0.99 and nT=0.90, respectively (Table S1). Based on these observations, we set hmSEEKR default transition parameters to be qT=0.99 and nT=0.90, enabling end-users to perform preliminary scans without extensive training. However, as best practice, end-users should conduct their own grid searches to define qT and nT to ensure transition parameters best capture the desired outputs of bespoke scans.

#### Additive searches with multiple query domains

Lastly, in our development and testing efforts, we encountered scenarios where we found it useful to search an RNA target(s) for similarity to multiple query domains in parallel. For example, aggregate similarity to multiple domains that harbor the same regulatory function may indicate an increased likelihood that the target RNA engages in that function. To this end, we created a summarizing search option in hmSEEKR, called “seqs-to-summary”, that simplifies parallel scans of target sequences against multiple query domains. Here, the same background sequences are used in all searches, and for each query in the list, end-users input individual qT and nT parameters. Once a “seqs-to-summary” search is complete, for each target sequence searched, outputs include: the number of hits to each query domain, the total number of nucleotides covered by each query, the median SEEKR-derived *p* value for all hits to each of the queries, and the total number of unique nucleotides and fraction of nucleotides in the target RNA that are covered by hits to the query domains being searched (Figure 1D).

### hmSEEKR successfully recovers query domains inserted randomly into transcriptomes

The lncRNAs *XIST*, *NEAT1*, and *MALAT1* are conserved, important for health, and expressed in the ENCODE Tier-1 K562 cell line in which enhanced crosslinking-immunoprecipitation (eCLIP) experiments have been performed (17, 18). They have also been subject to prior molecular genetic studies, including those that have mapped protein interactions and domains within each lncRNA that confer aspects of function (19–23). For these reasons, we focused our development and validation efforts on *XIST*, *NEAT1*, and *MALAT1*.

As an initial test of hmSEEKR’s search capabilities, we sought to determine whether the algorithm could successfully identify known domains after inserting the domains at random positions within a transcriptome. We separated *XIST*, *NEAT1*, and *MALAT1* into ∼1-2kb fragments using boundaries defined in part by prior deletion studies performed on each lncRNA (Figure 2A; (19–23)). We then inserted each fragment into 1000 different random locations within the set of expressed and chromatin-associated transcripts in K562 cells (24). This set of transcripts was defined using the probabilistic algorithm kallisto to align RNA-seq reads to a custom set of annotations that included all GENCODE v47 “comprehensive” transcript isoforms, and in addition, for each multiexonic gene, included a single “intron-containing” isoform that spanned the gene’s longest transcript annotation and contained all of its exons and introns (16, 25). We view these intron-containing isoforms as proxies for nascent transcripts produced from their host genes (26). We restricted our searches to chromatin-associated RNAs because our validation and discovery efforts centered around *XIST*, *NEAT1*, *MALAT1*, and other lncRNAs that have known or presumed cis-regulatory functions, all of which are thought to carry out their regulatory roles in the context of the chromatin environment. We limited our searches to expressed transcripts rather than searching the entire GENCODE comprehensive transcriptome, to enable cross-comparisons to eCLIP-defined protein interactions in subsequent validation efforts. Keeping with practice established for applications of the SEEKR algorithm, we tested hmSEEKR’s search capability at *k*-mer lengths *k* = 4, 5, and 6. We did not investigate *k*-mer lengths of *k* > 6. Because the length of the typical lncRNA domain is less than 4^6 nucleotides, quantifying similarity using *k*-mer lengths of *k* > 6 typically results in the majority of *k*-mers in the analyzed fragments having zero-count values, a scenario that can reduce SEEKR’s discriminatory power (6–8).

**Figure 2.**
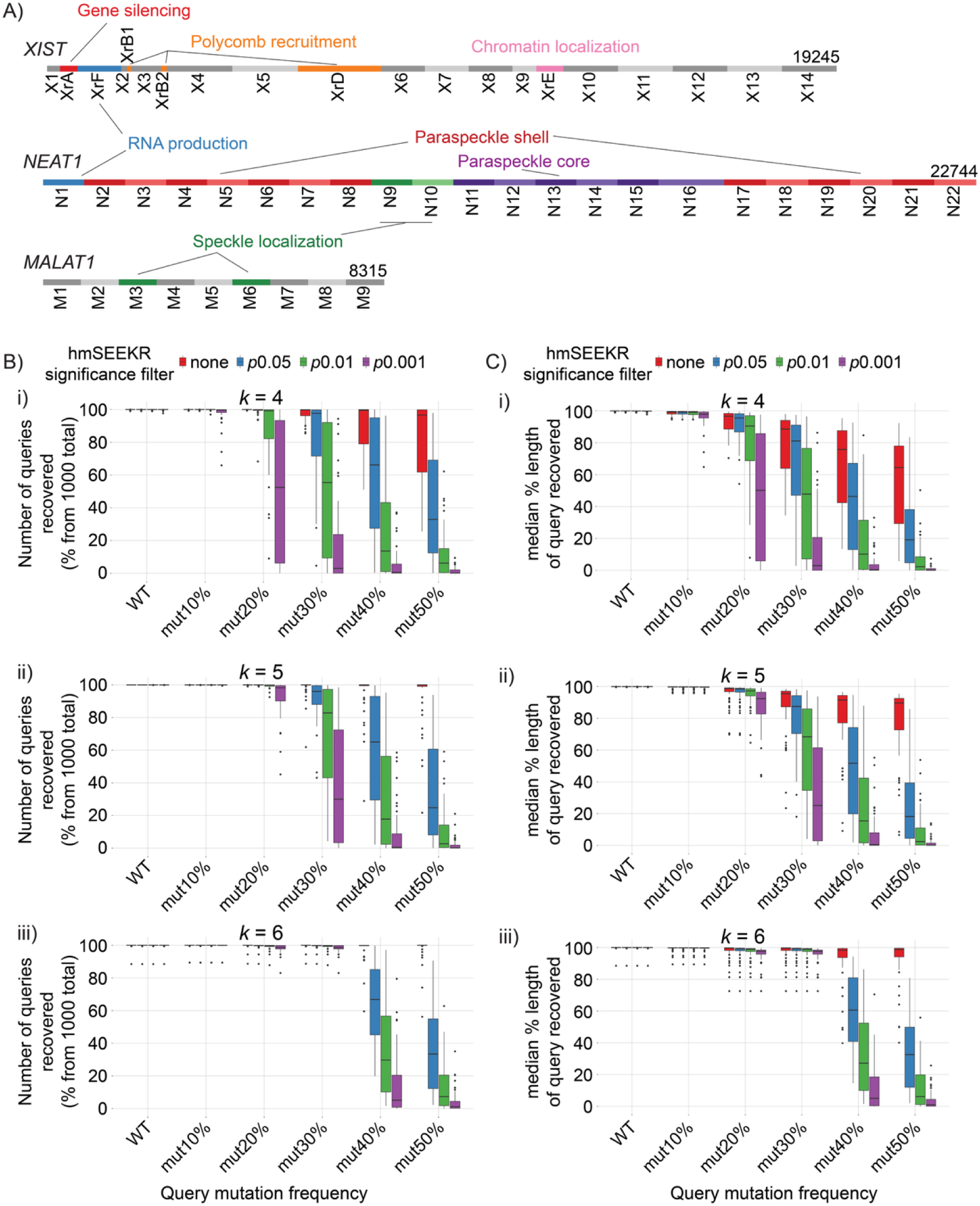
hmSEEKR successfully recovers query domains inserted randomly into transcriptomes. **(A)** Graphical overview of query domains defined in *XIST*, *NEAT1*, and *MALAT1* (*XNM*). Query partitions were established based on genetic deletion or complementation studies (18–23)**. (B)** Percent of *XNM* query insertions recovered (counts) and **(C)** median recovery of inserted *XNM* queries (by length/nucleotide) across increasing frequencies of query mutation, at *k*-mer length *k* = 4 (panel i), 5 (panel ii), or 6 (panel iii). At each mutational frequency tested, each *XNM* query was randomly inserted 1000 times into the expressed and chromatin-associated K562 transcriptome. See also Table S2.

We performed grid searches using GENCODE v47 lncRNAs as background to identify optimized transition parameters for each lncRNA fragment at *k*-mer lengths *k* = 4, 5, and 6, then used hmSEEKR to assess how well the algorithm recovered the randomly inserted query fragments from within the edited K562-cell transcriptomes, using a range of *p* value thresholds to retain hits (no threshold, *p* < 0.05, *p* < 0.01, and *p* < 0.001). At all thresholds and all lengths of *k*, hmSEEKR recovered 100% of all inserted fragments (20 *XIST*, 22 *NEAT1*, and 9 *MALAT1* parent fragments, counting partial recoveries of fragments), with a median recovery of ≥99.9% of the nucleotides across all fragments (Figures 2B,C; Table S2).

To determine how often hmSEEKR can recover different but related derivatives of the same fragments, we introduced increasing numbers of point mutations in each *XIST*, *NEAT1*, and *MALAT1* fragment, randomly inserted the mutated fragments into the K562 transcriptome, and measured the extent to which they could be recovered by hmSEEKR. We performed five tiers of analyses, in which we randomly mutated 10, 20, 30, 40, and 50 percent of the nucleotides in each inserted instance of each parent fragment, respectively. As the mutational frequency increased and the *p* value threshold for retaining hits decreased, hmSEEKR recovered fewer fragments and the median percent recovery of each fragment by length decreased, at all three *k*-mer lengths (Figure 2B, C). Of the *k*-mer lengths tested, *k* = 6 performed best, recovering a greater number of queries overall and on average, a greater percentage of each query at the higher mutational thresholds (Figure 2B, C; Table S2).

We conclude that the *k*-mer-based sequence search employed by hmSEEKR is sufficient to detect exemplar domains randomly inserted into sites across the human transcriptome, that varying the *p* value threshold for evaluating similarity between hit and query can be used to adjust the algorithm’s stringency of detection, and that among *k*-mer lengths *k* = 4, 5, and 6, *k* = 6 recovers the most inserted domains.

### hmSEEKR can recover individual and combinatorial RNA-protein interaction domains from within the transcriptome

Domains that confer function to lncRNAs often do so by interacting with specific sets of proteins (2). Thus, if hmSEEKR can identify hits within target RNAs that have the potential to carry out functions that are analogous to those of a query domain, we would hypothesize that the hits would interact with the same proteins as their corresponding query domains more than would be expected by random chance. To test this hypothesis, we searched the set of genes producing K562-expressed, chromatin-associated RNAs for hits to each of the same query fragments within *XIST*, *NEAT1*, and *MALAT1* as used in Figure 2A, at *k*-mer lengths *k* = 4, 5, and 6. We focused our analyses on genes producing chromatin-associated RNAs because *XIST*, *NEAT1*, and *MALAT1* are chromatin-associated themselves. We retained hits at SEEKR-derived *p* value thresholds of *p* < 0.05, *p* < 0.01, and *p* < 0.001 at all three *k*-mer lengths. As expected from our analyses in Figure 2, the number of hits to each query domain decreased as the *p* value threshold for hit definition increased in stringency at all three *k*-mer lengths (Figure 3A and Table S3).

**Figure 3.**
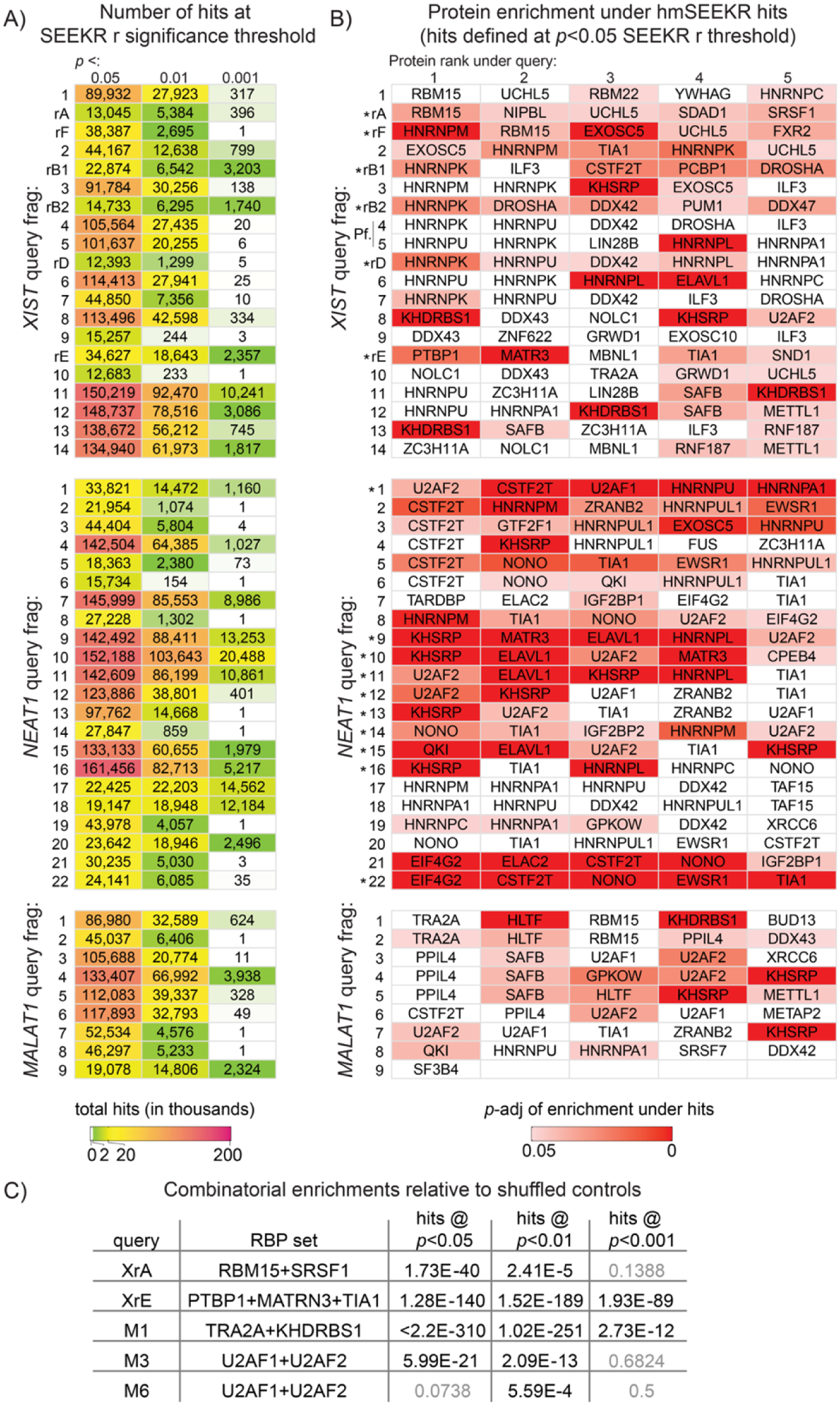
hmSEEKR can recover individual and combinatorial RNA-protein interaction domains from within the transcriptome. **(A)** Number of hits to each *XNM* query in the set of genes that produce chromatin-associated RNAs in K562 cells. **(B)** Significance of enrichment of CLIP signal under hmSEEKR hits versus shuffled controls for query-associated RBPs across all *XNM* query domains (Wilcoxon signed-rank test, Benjamini-Hochberg adjusted). *’s, query domains analyzed as part of comparison to minimal versions of *XIST* and *NEAT1* in Figure 5E. “Pf.”, PflMI fragment described in (20). **(C)** Significance of combinatorial enrichments of CLIP signal under hmSEEKR hits for *XIST* and *MALAT1* query domains that have been previously shown to interact with more than one protein client. Significance of combinatorial enrichment (Fisher’s exact) is shown at each of three separate *p* value thresholds for hmSEEKR hit definition. All searches performed at *k*-mer length *k* = 6. See also Figures S1-3 and Table S3.

Next, we sought to determine whether the same proteins that exhibit enriched association with query domains also exhibit enriched association over hits to the query domains. To approximate protein association in K562 cells, we used a set of previously collected eCLIP datasets measuring association with 139 different protein targets. For each query, we identified its top five associated proteins by ranking the difference of normalized reads per million aligned reads (RPM) over the query for each individual protein target of eCLIP minus the RPM for the same protein’s paired input control (hereafter referred to as “input-control normalized read densities”; (18). In parallel, for each set of hits to each query domain, we generated size-matched control regions by randomly shuffling the location of each hit once within the set of coordinates from genes producing chromatin-associated RNAs in K562 cells. Lastly, for each query, we determined whether its set of hits associated with any of the corresponding top-five proteins more than would be expected by random chance, by comparing input-control normalized read densities over hits to randomly shuffled control regions.

At the *p* < 0.05 threshold for hmSEEKR hit definition, hits from the majority of query domains in *XIST*, *NEAT1*, and *MALAT1*, respectively, exhibited associations that were significantly higher than randomly shuffled controls for at least one of their corresponding top five query-associated proteins, across all tested *k*-mer lengths (Figure 3B shows results from *k* = 6; Table S4 reports results from *k* = 4, 5, and 6; significance of enrichment determined by Benjamini-Hochberg corrected Wilcoxon rank test). The number of significant associations with hits over controls deceased as stringency of hit definition increased, consistent with the decreasing hit number at more stringent *p* value thresholds across all tested *k*-mer lengths (Figures S1, S2).

Reassuringly, for many query domains whose top interaction partners have been validated by orthogonal studies, we observed that their corresponding hits were also significantly enriched in interactions with the validated interaction partner. We would classify these interactions as higher confidence than those that have yet to be reproduced by orthogonal studies. This list includes: RBM15 and SRSF1 for hits to *XIST* Repeat A (27–29); HNRNPM for hits to *XIST* Repeat F (7, 28); HNRNPK for hits to *XIST* Repeats B and D (30, 31); PTBP1, MATR2, and TIA1 for hits to *XIST* Repeat E (32); NONO for hits to several fragments of *NEAT1* (22, 23, 33); CSTF2T for hits to *NEAT1* intervals 1 and 2 (34); KHDRBS1 for hits to *MALAT1* interval 1 (35, 36); and U2AF2 for hits to *MALAT1* intervals 3, 4, 6, and 7 (36). Likewise, additional paraspeckle-resident proteins present in the top five proteins of several *NEAT1* query domains (EWSR1, HNRNPA1, HNRNPM, and HNRNPUL1) were also significantly enriched in their interactions over hits against *NEAT1* queries (37).

At least two and three query domains in *XIST* and *MALAT1*, respectively, have been documented to interact with more than one protein client, possibly simultaneously (27–29, 32, 35, 36). We sought to determine whether hits to these domains also exhibited evidence for interactions with the same combinations of proteins. Because Wilcoxon rank test is only suitable to examine a single eCLIP dataset at a time, we sought to employ a categorical approach to assess significance of combined protein enrichments. We performed these assessments at *p* value thresholds for hmSEEKR hit definition of *p* < 0.05, *p* < 0.01, and *p* < 0.001 and across *k*-mer lengths *k* = 4, 5, and 6. Examining each query separately at each *p* value threshold, for every hit, we generated two randomly shuffled, size-matched control regions from within the set of gene coordinates that expressed chromatin-associated RNAs in K562 cells: a “satellite control” and an “anchor control”. The satellite control serves as a randomized proxy for the hit region, whereas the anchor control serves as the basis for comparison to both the hit and the satellite control. We calculated input-control normalized read densities for the proteins of interest within each hit and its respective anchor and satellite controls. We then counted the number of times that both proteins were enriched over hits relative to anchor controls versus the number of times that both proteins were not enriched. In parallel, to estimate background, we also counted the number of times that both proteins were enriched over the satellite relative to the anchor controls versus the number of times that both proteins were not enriched. We used the counts to populate a 2x2 contingency table, and used Fisher’s exact test to evaluate whether the differences between hit-versus-anchor and satellite-versus-anchor were significant. At *k*-mer length *k* = 6, at the *p* < 0.05, *p* < 0.01, and *p* < 0.001 thresholds, hmSEEKR identified hits that exhibited a higher frequency of combinatorial interactions than expected from satellite and anchor controls in 4/5, in 5/5, and in 2/5 instances, respectively (Fisher’s exact; Figure 3C). Similar results were obtained a *k*-mer length *k* = 4 (combinatorial significance in 3/5, 4/5, and 2/5 instances) and at *k* = 5 (combinatorial significance in 4/5, 4/5, and 1/5 instances; Figure S3).

We conclude that using individual domains as queries, hmSEEKR can identify hits within the transcriptome that exhibit significant individual and combinatorial interactions with sets of proteins that also exhibit interactions with the corresponding queries. Moreover, across our validation efforts in Figures 2 and 3, we observed that *k*-mer length *k* = 6 tended to outperform *k*-mer lengths *k* = 4 and 5 (albeit, typically by only modest amounts). For that reason, in our applications below, we use *k*-mer length *k* = 6 as a primary search metric.

### hmSEEKR identifies *XIST*-, *NEAT1*-, and *MALAT1*-like elements in protein-coding and lncRNA genes

Having determined that hmSEEKR can successfully identify RNA elements that have similar *k*-mer contents and interact with similar sets of proteins as their corresponding query domains, we began to investigate examples of biological applications that hmSEEKR might enable. We first sought to quantify the transcriptome-level similarity to *XIST*, *NEAT1*, and *MALAT1* in greater detail. Within mammals, *XIST*, *NEAT1*, and *MALAT1* are among the most conserved and abundant nuclear lncRNAs, and while they are likely to be unique in many respects, several of the proteins that they interact with are present at near-micromolar concentrations in the cell and presumably interact with thousands of other RNAs (2, 38). Thus, the protein-interaction domains within *XIST*, *NEAT1*, and *MALAT1* likely share features with regions in other RNAs. Our analyses in Figure 3A support this idea, showing that under our least stringent search threshold of *p* < 0.05, thousands of *XIST*-, *NEAT1*-, and *MALAT1*-like elements were distributed amongst the set of K562-expressed, chromatin-associated RNAs. At the same time, as search stringency increased to *p* < 0.001, the number of *XIST*-, *NEAT1*-, and *MALAT1*-similar regions dropped, precipitously in many instances, down to 200 or fewer hits in as many genes that produce K562-expressed, chromatin-associated RNAs (Figures 3A, 4A). For example, following searches performed with *k*-mer length *k* = 6, nine of 20 query domains within *XIST*, 10 of 22 query domains within *NEAT1*, and five of nine query domains in *MALAT1* registered hits within < 200 genes at the *p* < 0.001 threshold (Figure 4A). Other domains registered hits within > 500 genes even at the *p* < 0.001 threshold, including Repeat B, Repeat E, and query domains #11-14 in *XIST*, fragments #9-11 in *NEAT1*, and fragments #4 and 9 in *MALAT1* (Figure 4A).

**Figure 4.**
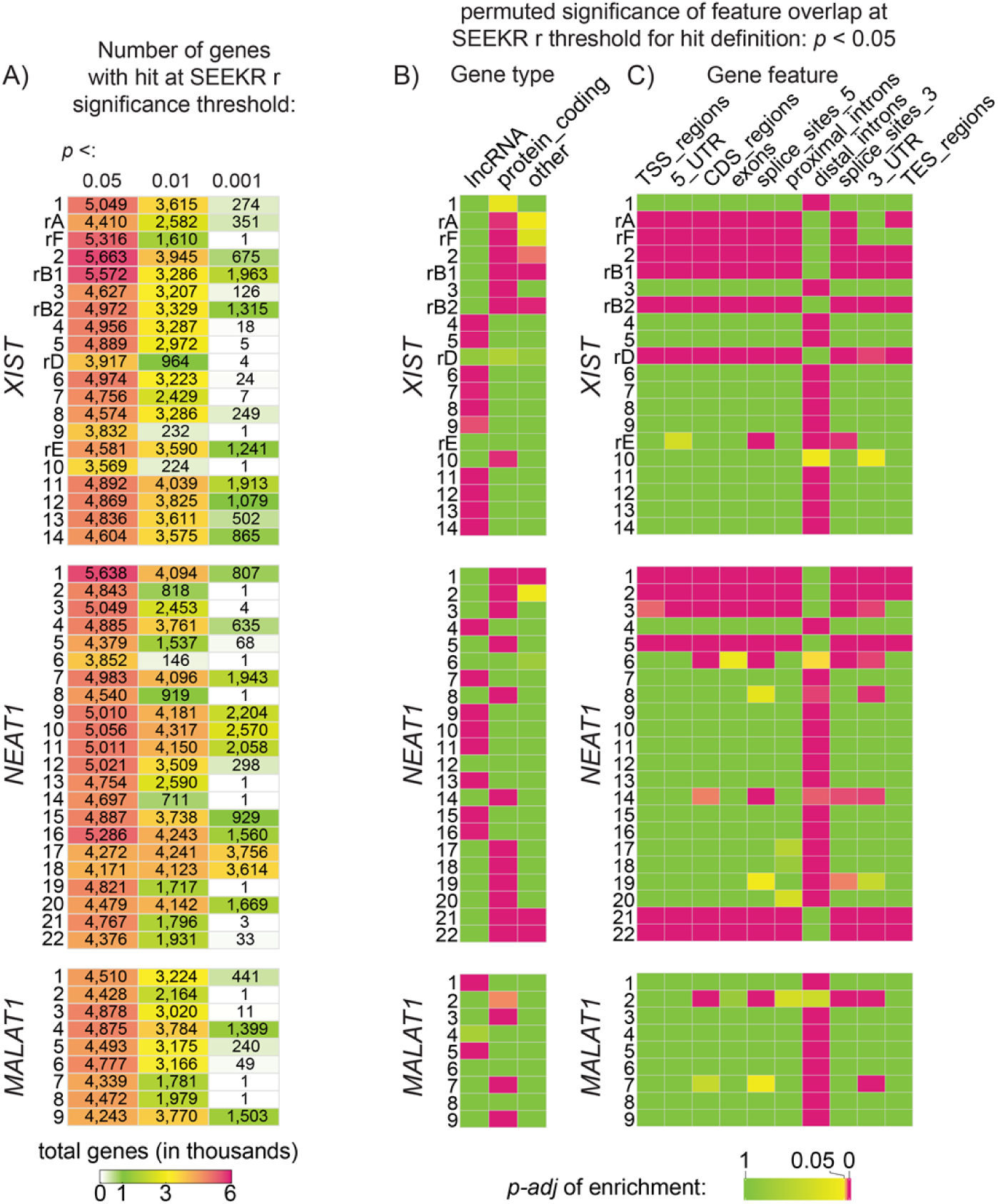
hmSEEKR identifies *XIST*-, *NEAT1*-, and *MALAT1*-like elements in protein-coding and lncRNA genes. **(A)** Number of genes producing chromatin-associated RNAs in K562 cells that contain hits to *XNM* queries at different *p* value thresholds of hit definition. **(B)** For each *XNM* query, adjusted *p* value of hit enrichment in the set of genes that produce chromatin-associated RNAs in K562 cells (Fisher’s exact, Benjamini-Hochberg adjusted); and **(C)** adjusted *p* value of hit enrichment in the features of genes that produce chromatin-associated RNAs in K562 cells (Fisher’s exact, Benjamini-Hochberg adjusted). Hits were defined using a SEEKR-derived *p* value threshold of p < 0.05. All searches performed at *k*-mer length *k* = 6. See also Figure S4 and Table S3.

Given the detection of *XIST*-, *NEAT1*-, and *MALAT1*-like elements throughout the transcriptome, we next investigated whether the elements preferentially overlapped with certain gene types or gene features. For each *XIST*-, *NEAT1*-, and *MALAT1*-like element detected, we randomly shuffled its location among genes producing K562-expressed, chromatin-enriched RNAs, then determined the proportion of real and shuffled hits that overlapped with the GENCODE-annotated gene types and transcript features listed in Figures 4B and C. We used Fisher’s exact test followed by Benjamini-Hochberg correction to determine whether hits to each query domain were significantly enriched in gene types or features relative to shuffled controls. At the *p-adj.* < 0.05 threshold, we found that hits to query domains in all three transcripts were enriched within both lncRNA and protein-coding genes (Figure 4B). The enrichments in hits to *XIST*, *NEAT*, and *MALAT1* query domains were split between lncRNA and protein-coding genes: hits to 10 and seven *XIST* domains were enriched in lncRNA and protein-coding genes, respectively; hits to eight and 12 *NEAT1* domains were enriched in lncRNA and protein-coding genes, respectively; and hits to two and four *MALAT1* domains were enriched in lncRNA and protein-coding genes, respectively (Figure 4B).

Turning next to gene features, we found that the majority of hits against queries from all three lncRNAs were enriched in distal introns (intronic sequence > 2000 nt away from the nearest exon; Figure 4C). Conversely, queries whose hits were not enriched in distal introns tended to be enriched across all other genic features, for example: hits to XIST Repeats A, F, B, and D and hits to domains at the 5′ and 3′ ends of *NEAT1* (in its RNA production and paraspeckle shell regions (22, 23); Figure 4C). With the exception of its intervals 2 and 7, hits to *MALAT1* were exclusively enriched in distal introns (Figure 4C). The trend of enrichment towards distal introns was observed at the *p* < 0.01 and *p* < 0.001 thresholds, as well (Figure S4).

Thus, despite their identity as lncRNAs, we observed that query domains across *XIST*, *NEAT1*, and *MALAT1* were similar to regions contained within transcripts produced from both lncRNA and protein-coding genes. Hit regions were enriched in many genic features, most frequently within distal introns. While certain query domains across *XIST*, *NEAT1*, and *MALAT1* had significant regional similarity to thousands of other transcript regions, other domains were similar to far fewer regions and thus appeared more unique within each lncRNA.

### Sequential hmSEEKR searches identify K562-expressed transcripts that resemble minimal versions of *XIST* and *NEAT1*

Having found that hmSEEKR can detect individual regions of similarity to *XIST*, *NEAT1*, and *MALAT1*, we next sought to determine whether any expressed transcripts exhibited evidence of sequential, domain-level similarity to the lncRNAs. Sequential, domain-level similarities might be more likely to suggest functional similarities than similarities to individual domains. For this analysis, we focused on *XIST* and *NEAT1*, because genetic deletion and reconstruction experiments have identified domains within them that are essential for their functions and comprise minimally functional versions of each lncRNA. Although full-scale silencing by *XIST* requires the entirety of its sequence, its Repeat A, F, B1/2 and D, and E domains are critical for its ability to silence gene expression, produce high levels of its own transcript, recruit Polycomb complexes, and remain tethered near the inactive X chromosome, respectively, and can be considered as components of a minimal *XIST* ((39); Figure 2A). Similarly, the combination of domains #1, #9-16, and #22 are sufficient for *NEAT1* to form an organized, paraspeckle-like structure in human cells ((22); Figure 2A). We therefore sought to examine whether any K562-expressed, chromatin-associated RNAs contained regional similarities that were present in sequential patterns resembling those found in the minimal functional versions of *XIST* or *NEAT1*. Such RNAs, if they existed, might represent candidates for carrying out *XIST*- or *NEAT1*-like functions in cells. Limiting our searches to K562-expressed and chromatin-associated RNAs enabled us to use eCLIP data as a way to cross-reference our search results with protein interactions in the RNAs identified.

Focusing first on *XIST*, using *k*-mer length *k* = 6, we searched the set of K562-expressed, chromatin-associated RNAs for transcripts that resembled a minimal functional version of *XIST*. To be classified as *XIST*-similar in this search, we required transcripts to contain a hit to Repeats A and F within their first 3kb, followed by hits to both Repeats B1/2 and D, followed by a hit to Repeat E (39). We acknowledge that these restrictions have the potential to exclude bona fide *XIST*-similar transcripts whose internal domain structure does not meet the restrictions above. However, because our primary goal was to test a use-case of hmSEEKR rather than perform an exhaustive, experimentally validated search of *XIST* similarity, we required restrictions to be in place.

At the p < 0.05 threshold, we detected 674 *XIST*-similar transcripts, including the nascent and spliced annotations of *XIST* itself (Figure 5A; Table S5). All but seven *XIST*-similar transcripts at the *p* < 0.05 level (99%) were either intron-containing (i.e. nascent) or intronless/monoexonic annotations. 626 (93%) and 45 (7%) of the 674 *XIST*-similar transcripts originated from protein-coding and lncRNA genes, respectively (and an additional three transcripts were classified as “transcribed_unprocessed_pseudogenes”; Table S5). Among the *XIST*-similar lncRNAs were many that remain largely unstudied as well as several that have been implicated in aspects of development or disease, including *MIR17HG*, *SNHG17*, and the *XIST*-adjacent lncRNA *JPX* (Table S5). *KCNQ1OT1*, a known repressive lncRNA that has previously been shown to contain *XIST*-like sequence features (7, 13), was not identified here owing to its lack of detectable Repeat A-like domain within the first 3kb of its primary transcript. *KCNQ1OT1* did, however, meet the other criteria to be defined as *XIST*-similar, and it met our complete criteria to be classified as *NEAT1*-similar in searches below. Intriguingly, the nascent annotations of several genes producing RNA-binding proteins that associate with or directly bind *XIST* were also identified as *XIST*-similar, including *ACIN1*, *CIZ1*, *CPSF6*, *CPSF7*, *HNRNPA2B1*, *HNRNPC*, *HNRNPK*, *HNRNPM*, *HNRNPU*, *LBR*, *LUC7L2*, *LUC7L3*, *MATR3*, *SPEN*, and *ZC3H18* ((40) and references therein). As stringency of hit definition moved to *p* < 0.01, the only transcripts detected as *XIST*-similar, other than *XIST* itself, were the nascent annotations of two protein coding genes, *CSPP1* and *LDAH*; at the *p* < 0.001 threshold, other than *XIST*, no transcripts were identified as *XIST*-similar (Table S5).

**Figure 5.**
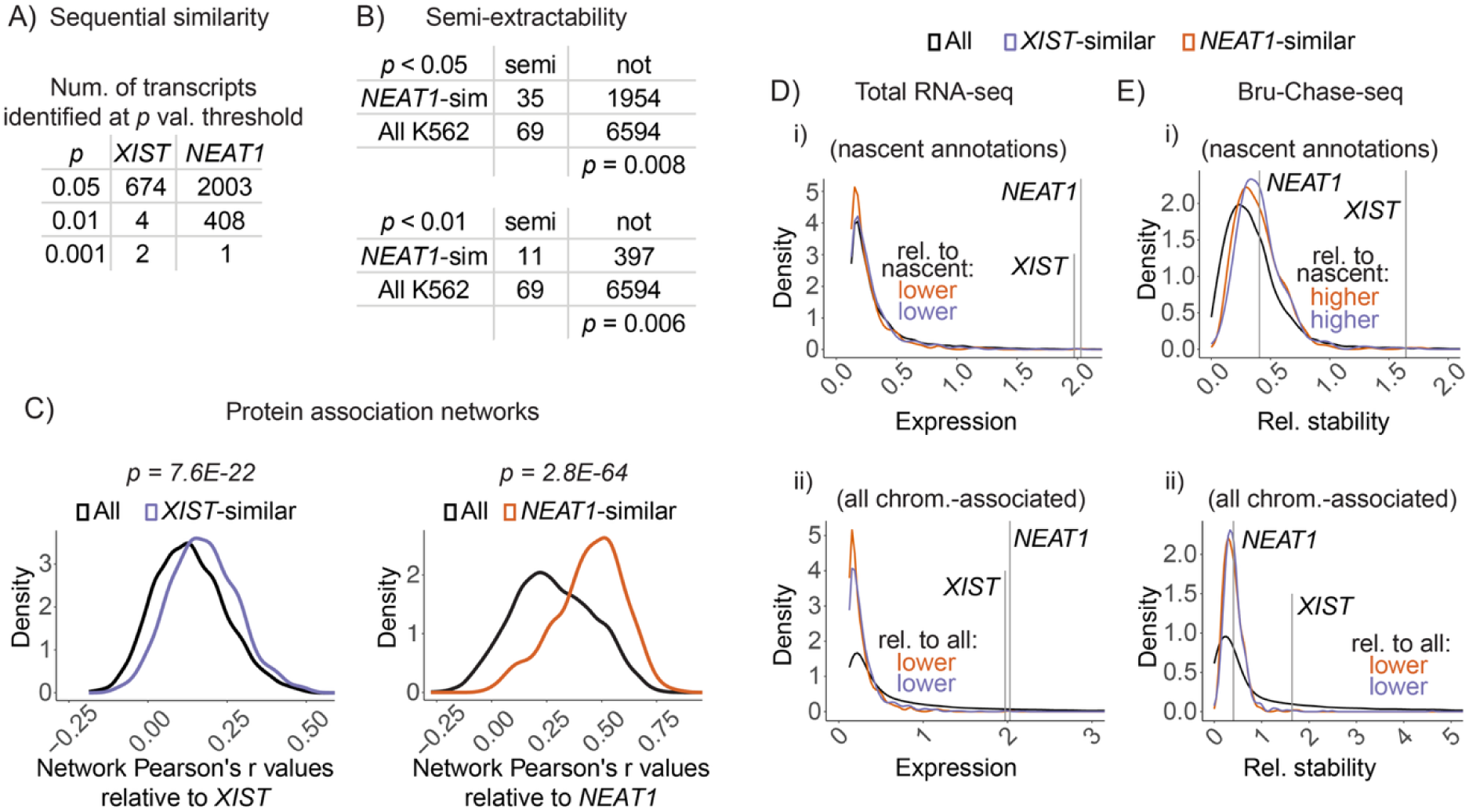
Sequential hmSEEKR searches identify K562-expressed transcripts that resemble minimal versions of *XIST* and *NEAT1*. **(A)** Numbers of *XIST*- and *NEAT1*-similar transcripts at different *p* value thresholds of hmSEEKR hit definition. **(B)** Fisher’s exact contingency tables comparing K562-expressed semi-extractable vs normally-extractable RNAs, at *p* value thresholds of hmSEEKR hit definition of *p* < 0.05 and *p* < 0.01. Only NEAT1 itself was detected at *p* < 0.001. **(C)** Protein association networks of *XIST*- and *NEAT1*-similar genes compared to networks from all genes producing K562-expressed, chromatin-associated RNAs. *p*, Wilcoxon-rank sum tests. **(D, E)** Expression quantified by total RNA-seq (D) and relative stability quantified by Bru-Chase-seq (E) for *XIST*- and *NEAT1*-similar transcripts relative to all transcripts, using either **(i)** only intron-containing/nascent annotations or **(ii)** all K562-expressed, chromatin-associated transcripts including spliced annotations. Whether the distribution of *XIST*-and *NEAT1*-similar transcripts was significantly higher or lower than the distribution of all transcripts compared is indicated on each plot, with text colored by its corresponding distribution (Wilcoxon rank-sum). All searches performed at *k*-mer length *k* = 6. See also Tables S5 and S6.

Turning next to minimal *NEAT1* (22), we searched the set of K562-expressed, chromatin-associated RNAs for transcripts that contained sequential hits to domain #1 within their first 3kb, followed by hits to domains #9 or 10, followed by at least 1kb of hits to any domain from #11-16, and lastly a hit to domain #22. As with our *XIST*-based searches above, we acknowledge that these restrictions have the potential to exclude bona fide *NEAT1*-similar transcripts whose internal organization differs from minimal *NEAT1* defined in (22). In total, we detected 2003, 408, and 1 *NEAT1*-similar transcripts at the *p* < 0.05, 0.01, and 0.001 thresholds, respectively (Figure 5A; Table S5). At the *p* < 0.05 threshold, 93 and 6 percent of *NEAT1*-similar transcripts arose from protein-coding and lncRNA genes, respectively, and 99 percent were intron-containing. At the *p* < 0.01 threshold, 93 and 7 percent of *NEAT1*-similar transcripts arose from protein-coding and lncRNA genes, respectively, and 100 percent were intron-containing. At the *p* < 0.001 threshold, other than *NEAT1*, no transcripts were identified as *NEAT1*-similar (Table S5). The lncRNA *MALAT1* was not detected as *NEAT1*-similar, supporting the view that *NEAT1* and *MALAT1* have diverged in function since their proposed evolutionary origin from a gene duplication event in ancient tetrapods (41).

*NEAT1* (but not *XIST*) has been found to exhibit the unusual property of being “semi-extractable” during RNA extraction methods that rely on “tri-reagent”, mixtures of acid guanidinium thiocyanate, phenol, and chloroform that separate RNA from proteins and DNA (42). Rather than partitioning into the aqueous phase after cell-solubilization with tri-reagent, *NEAT1* predominantly partitions into the proteinaceous interphase, unless heat or needle shearing are applied; hence its classification as semi-extractable (42). Premising that *NEAT1*-similar transcripts might share properties with *NEAT1*, we examined whether *NEAT1*-similar transcripts exhibited evidence of semi-extractability. Semi-extractable RNAs produced from a total of 1074 genes were recently identified across five human cell lines (43), and 69 of these genes produced transcripts present in our set of K562-expressed, chromatin-associated RNAs. Limiting our comparisons to intron-containing or intronless transcripts (because nearly all *NEAT1*-similar transcripts were intron-containing or intronless), we found that *NEAT1*-similar transcripts at the *p* < 0.05 and *p* < 0.01 thresholds were significantly more likely to be semi-extractable than expected by chance, providing additional evidence that our sequential search strategy can identify RNAs that share biological properties (Figure 5B; Fisher’s exact; Table S5 (42, 43)).

As a separate way to connect our sequential search results to biological properties of the RNAs identified, we examined patterns of protein association as measured by eCLIP. If the sequential domain similarities detected by hmSEEKR were biologically meaningful, we might expect *XIST*- and *NEAT1*-similar transcripts to exhibit patterns of protein association that more closely resembled those of their corresponding query lncRNAs than other RNAs. Indeed, in a prior study performed in mouse trophoblast stem cells, we found that the patterns of protein association within the three known repressive lncRNAs *Xist*, *Airn*, and *Kcnq1ot1* were more similar to each other than to the majority of other chromatin-associated transcripts, implying that the similar patterns reflected similarities in lncRNA regulatory function (14).

We focused our protein association analyses on transcripts detected at the *p* < 0.05 threshold for *XIST* similarity, because more stringent thresholds failed to identify significant numbers of *XIST*-similar transcripts other than *XIST*, and at the *p* < 0.01 levels for *NEAT1* similarity (rather than *p* < 0.05), because we identified hundreds of RNAs at this threshold and wished to restrict our analyses to the set of RNAs exhibiting stronger levels of similarity to *NEAT1* (Figure 5A). We likewise restricted our analyses to gene bodies (i.e. intron-containing and intronless annotations), which enabled us to use eCLIP genomic alignments without having to introduce ambiguity related to spliceforms, and simplified our comparisons in a way that was justified given that 99 and 100 percent of *XIST*- and *NEAT1*-similar RNAs were intron-containing or intronless. In total, this filter retained the genic coordinates of 667 and 408 *XIST*- and *NEAT1*-similar transcripts, respectively.

To define patterns of protein association across our genes of interest, rather than examine all 139 eCLIP datasets, we assembled aggregate lists of the top five proteins most-enriched by eCLIP within the query domains that comprised minimal *XIST* and *NEAT1* (rows marked by asterisks in Figure 3B). We further restricted the lists to retain only those proteins whose eCLIP read density was significantly higher under hmSEEKR hits compared to randomly shuffled controls (shaded boxes in rows marked by asterisks in Figure 3B). These restrictions limited our analyses to the proteins with the most robust eCLIP signal over our query domains of interest and which exhibited evidence of sequence-driven associations in other RNAs. Reassuringly, the restrictions retained most if not all of the known protein co-factors of each query domain that were profiled by eCLIP (e.g. RBM15, SRSF1, HNRNPK, and PTBP1, MATR3, and TIA1 in *XIST*, and many paraspeckle-resident proteins in *NEAT1*). At the same time, the restrictions likely reduced the inclusion of functionally irrelevant proteins in our lists. In total, we retained 21 and 17 eCLIP’d proteins in our *XIST* and *NEAT1* lists, respectively.

With these lists, we sought to establish networks that quantify the patterns of protein association within each gene being evaluated; in turn, these networks provide a length-independent metric that can be used to quantify patterns of protein association between *XIST*-and *NEAT1*-similar genes and their respective query lncRNAs (14). To this end, for each protein in each list, we generated background-corrected counts in 25 nt bins across each gene producing a chromatin-associated RNA in K562 cells. Networks of protein associations were defined for each gene by performing Pearson’s correlations between background-corrected, binned eCLIP data for all possible pairs of proteins. We then compared the protein association network of each gene to the analogous network in its respective query lncRNA, either *XIST* or *NEAT1*, again using Pearson’s correlation. Lastly, we used Wilcoxon rank-sum test to compare the distribution of Pearson’s r values between *XIST*- and *NEAT1*-similar genes and their corresponding query lncRNAs (*XIST* or *NEAT1*) versus the distribution of Pearson’s r values between all genes producing a chromatin-associated RNA in K562 cells and the corresponding query lncRNAs (again, *XIST* or *NEAT1*). Strikingly, relative to all genes, the protein association networks for *XIST*-and *NEAT1*-similar genes were significantly more similar to the networks of *XIST* and *NEAT1*, respectively, than would be expected by chance (*p* = 7.6E-22 and 2.8E-64 for the *XIST*- and *NEAT1*-similar distributions, respectively; Wilcoxon rank-sum; Figure 5C; Table S5).

Intuitively, the extent of gene regulatory function exerted by a lncRNA might be a product of its internal domain content, its abundance, and its stability. For example, the lncRNA *XIST/Xist* is more stable and more abundant than many other lncRNAs (13), and its ability to regulate gene expression is exceptional (*XIST/Xist* represses an entire chromosome). We therefore quantified the abundance and stability of *XIST*- and *NEAT1*-similar transcripts relative to all K562-expressed and chromatin-associated RNAs, examining total RNA- and Bru-Chase-seq data from ENCODE (24), reasoning that the more abundant and more stable the RNA, the more likely it is to exert a potent biological effect. We found that on average, intron-containing/nascent *XIST*- and *NEAT1*-similar transcripts were significantly lower in abundance but significantly more stable than other intron-containing/nascent isoforms classified as K562-expressed and chromatin-associated RNAs (*p* < 0.0002 in all cases; Wilcoxon rank-sum; Figure 5Di, Ei). Expanding our analyses to compare to all K562-expressed and chromatin-associated transcripts including spliced isoforms, we found that *XIST*- and *NEAT1*-similar transcripts were significantly lower in abundance and less stable when (*XIST*-similar vs. all chromatin-associated, *p* < 1E-10 in all cases; Wilcoxon rank-sum; Figure 5Dii, Eii). Of all *XIST*-similar transcripts, *XIST* itself was the sixth most stable and fourth most abundant (Figures 5D and 5E; Table S5). In contrast, of all *NEAT1*-similar transcripts, *NEAT1* ranked 167th in stability (out of 408 transcripts) and second in abundance, behind the intron-containing annotation of the protein-coding gene *PRAME* (Figures 5D and 5E; Table S5).

Lastly, we sought to determine the extent to which *XIST*- and *NEAT1*-similar RNAs exhibited evidence consistent with being engaged in the local regulation of gene expression. *XIST* is the most repressive lncRNA known, functioning to silence transcription over one entire X chromosome in cis (i.e. on the same X chromosome from which it is expressed; (15)). In contrast, *NEAT1* is an architectural RNA whose effects on gene expression are complex and predominantly occur in trans; although in knockout mice, there is evidence that *NEAT1* mediates both local cis repressive and activating effects (44–46). Thus, under a simplistic hypothesis, we might expect RNAs that exhibit *XIST*-like properties to be more likely to exhibit evidence of repressing nearby genes than RNAs that exhibit *NEAT1*-like properties.

To test this hypothesis, we examined RNA-seq data from the Genotype-Tissue Expression (GTEx) project (47). Specifically, across a subset of GTEx tissues, we retained genes that were differentially expressed, in order to focus our analyses on genes whose expression was variable across tissues. Then, for each retained *XIST*- and *NEAT1*-similar gene, we examined the extent and manner in which its expression correlated with the expression of other retained genes nearby; “nearby” being defined as located within 1 megabase surrounding the transcriptional start site of the parent *XIST*- or *NEAT1*-similar gene. From these analyses, we found that as a class, *XIST*-similar genes were no more likely than *NEAT1*-similar genes to exhibit a greater number of negative correlations with nearby genes than positive ones; 93 versus 360 retained *XIST*-similar genes and 62 versus 248 retained *NEAT1*-similar genes exhibited more negative correlations with nearby genes than positive ones (p = 0.93, Fisher’s exact test; Table S6). Thus, the simplistic expectation – that *XIST*-similar genes would more frequently exhibit negative correlations with nearby genes than *NEAT1*-similar genes – was not met.

However, on a case-by-case basis, these analyses highlighted several *XIST*- and *NEAT1*-similar genes whose expression across GTEx tissues exhibited strong enrichments for negatively or positively correlated nearby genes. Although causality cannot be established from these correlations – effects driven by DNA regulatory elements seem just as likely or more likely to dominate than effects driven by putative regulatory RNAs – our analyses highlight *XIST*- and *NEAT1*-similar RNAs that are located in what appear to be coordinately regulated genomic regions and which might represent priority targets for future study.

Overall, we found that *NEAT1*-similar RNAs were more frequently located near positively and negatively correlated genes than *XIST*-similar RNAs. The median number of positively and negatively correlated genes within 1 megabase of *NEAT1*-similar RNAs were 6 and 2, respectively. In contrast, the median number of positively and negatively correlated genes within 1 megabase of *XIST*-similar RNAs were 3 and 1, respectively. The upper quartiles of positively and negatively correlated genes within 1 megabase of *NEAT1*-similar RNAs were 10 and 4, respectively, and the same numbers for *XIST*-similar RNAs were 6 and 3, respectively (Table S6). While any of single correlation we observed has the potential to be RNA-influenced, most intriguing to us were the genes producing *XIST*- and *NEAT1*-similar RNAs that exhibited the strongest biases in the number of nearby positively or negatively correlated genes. Minimally, we can conclude that these genes have the potential to produce RNAs that engage in *XIST*- or *NEAT1*-similar functions and sit in genomic regions that exhibit strong signatures of directionally-biased, coordinate regulation. We define a “correlation bias” as the number of negatively correlated genes subtracted from the number of positively correlated genes; genes with positive correlation biases are surrounded by more positively correlated genes than negatively correlated ones, and genes with negative correlation biases are surrounded by more negatively correlated genes than positively correlated ones. Twenty-one *XIST*-similar RNAs had negative correlation biases of 5 or more, and four *XIST*-similar RNAs had negative correlation biases of 12 or more. In contrast, 121 *XIST*-similar RNAs had positive correlation biases of 5 or more, and 26 *XIST*-similar RNAs had positive correlation biases of 12 or more. We observed similar trends for *NEAT1*-similar RNAs. Twenty-three *NEAT1*-similar RNAs had negative correlation biases of 5 or more, and two had negative correlation biases of 12 or more; and 138 *NEAT1*-similar RNAs had positive correlation biases of 5 or more, while 36 had positive correlation biases of 12 or more. Notably, *NEAT1* itself exhibited a negative correlation bias of 5, harboring 6 negatively and one positively correlated gene in its genomic surroundings; three of these genes were previously identified as being dysregulated in the mouse brain upon *Neat1* knockout, suggesting that *NEAT1* may exhibit conserved cis-regulatory effects ((44); Table S6).

In summary, examining RNAs with combinatorial, sequentially-arrayed hits to a minimal version of *XIST*, we identified 674 *XIST*-similar transcripts at the *p* < 0.05 threshold, most of which originated from protein-coding genes, and nearly all of which were less abundant and less stable than *XIST*. At a more stringent search threshold (*p* < 0.01), we were able to identify only two other *XIST*-similar transcripts, both of which were nascent annotations produced from protein-coding genes. At the *p* < 0.01 threshold, we identified hundreds of *NEAT1*-similar transcripts, including several transcripts previously found to share the property of semi-extractability (43), and at least one, the intron-containing/nascent transcript from the protein-coding gene PRAME, that kallisto alignment estimated as being more abundant than *NEAT1*. Overall, the protein association networks of *XIST*- and *NEAT1*-similar transcripts were significantly more similar to the networks within their query lncRNAs than would be expected by chance, and many of the transcripts are located within clusters of coordinately regulated genes, linking our sequence-based search results to orthogonal experimental observations and nominating potential regulatory relationships for future study.

### hmSEEKR identifies lncRNAs that harbor local similarities to known regulatory domains

Our observation that combinatorial hmSEEKR scans can identify target RNAs whose networks of protein interaction resembled those of their query lncRNAs support the view that sequential, domain-based hmSEEKR scans can identify biologically meaningful similarities between RNAs. We next sought to expand our investigations to lncRNAs that have previously been linked to cis-regulatory functions, to determine whether domain-based scans might be leveraged to provide testable insights.

As a mechanism of transcriptional activation, several presumed *cis*-activating lncRNAs have been proposed to directly bind protein effector complexes, including the Mediator, WDR5/MLL, and CBP/p300 complexes (48–50). At the same time, it has been observed that *Xist/XIST* and other cis-repressive lncRNAs interact with effector complexes through bridged interactions with RNA-binding proteins, rather than by direct binding (13, 14, 30, 51, 52). Perhaps consistent with those observations, direct RNA binding by WDR5/MLL has recently been shown to inhibit its enzymatic activity (53). Still other lncRNAs are thought to influence gene expression through RNA sequence-independent mechanisms that involve promoter elements or the act of transcription (54, 55). The extents to which lncRNAs engage with effector complexes through direct or protein-bridged interactions, or influence transcription via sequence-independent mechanisms, remain unclear. Accordingly, across cis-regulatory RNAs, it remains unclear whether shared RNA sequence features underlie mechanistically relevant RNA-protein interactions, and whether recurring features are preferentially enriched within activating or repressive lncRNAs. In most cases, it is even unclear whether the nascent lncRNA or its spliced lncRNA product is the relevant regulatory molecule. Given the unknowns, we reasoned that an application of hmSEEKR might provide insights, by quantifying the extent of similarity between RNA domains that have previously linked to discrete molecular functions within a cohort of cis-acting regulatory RNAs.

To this end, we assembled a collection of 69 query domains with known or inferred molecular functions from within the lncRNAs *XIST*, *NEAT1*, and *MALAT1*, along with domains in other lncRNAs as well as enhancer RNAs (eRNAs) that collectively have been implicated in nuclear export, nuclear retention, chromatin retention, and transcriptional activation (Table S7; (19–23, 27, 49, 50, 55–62)). We also curated a list of known or putative regulatory lncRNAs whose expression has been tied to transcriptional activation or repression in cis. In total, we selected 36 and 18 lncRNAs previously linked to cis-activating and cis-repressive functions, respectively (Table S8; (13, 19, 20, 48, 49, 63–105)). Because lncRNAs are often spliced inefficiently (106–109), and because *XIST*, *NEAT1*, and *MALAT1* exhibit significant regional similarity to intronic sequences (Figure 4), we retained one representative intron-excluding (i.e. spliced) and one representative intron-including (i.e. nascent) isoform for each lncRNA gene in our searches. This strategy enabled us to determine whether the spliced or nascent transcript, or both, harbors similarity to functional domains in our list. Of note, the locus producing the lncRNA *PVT1* has been shown to exert both activating and repressive effects depending on the context studied; the activating effects mediated possibly through its nascent transcript or internal enhancers (103, 110–113), and the repressive effects mediated through specific spliced isoforms or its promoter (114–117). In our analyses below, using the MANE annotation, we classified the nascent transcript produced from the *PVT1* locus as an activator, and its spliced transcript as a repressor (116, 118).

We then searched our sets of known or presumed activating and repressive lncRNAs for similarity to our collection of query domains, to examine whether hits to queries exhibited a bias within one set of lncRNAs over the other. We first performed grid searches to identify optimal hmSEEKR search parameters for each query domain, then scanned our collection of regulatory lncRNAs, retaining hits at the *p* < 0.05, 0.01, and 0.001 thresholds, using transition parameters and length specifications listed in Table S1. We adopted the following strategy to assess whether query domains exhibited hits that were preferentially enriched within our sets of presumed activating or repressive RNAs, calculating enrichments separately for sets of intron-excluding (i.e. spliced) and intron-including (i.e. nascent) isoforms. For each query, we calculated the summed length of its hits within each set of lncRNA isoforms examined. We then used Fisher’s exact test to determine whether the summed hit lengths were significantly enriched within the sets of activating versus repressive lncRNAs. Prior to performing Fisher’s exact, we scaled all hits and lncRNA isoforms by their lengths rounded to the nearest hundred nucleotides, which reduced counts to a level that required a greater difference in overall nucleotide length to ascribe significance. By these analyses, considering all *p* value thresholds for hmSEEKR hit definition, we observed 13 and 36 query domains that exhibited a hit enrichment within either the sets of presumed activating or repressive lncRNAs, respectively. An additional 10 query domains exhibited enrichment in both the activating and repressive sets of lncRNAs, but within the spliced or nascent class in one set and the opposite class in the other, or within the activating or repressive set of lncRNAs at one hmSEEKR search threshold and the opposite class in another (Table S9).

Query domains whose hits were exclusively enriched towards presumed activating lncRNAs included the MLL-associated region within *HOTTIP* (49), proposed P300-recruiting eRNAs produced from enhancers near the genes *Sp3* and *YY-1* (50), and the *ncRNA-a7/TRERNA* eRNA (48). Query domains whose hits were exclusively enriched towards presumed repressive lncRNAs included four of the five of the query fragments from a heterogeneous nuclear ribonucleoprotein (HNRNP)-binding domain of *ASAR6-141*, a lncRNA required for proper replication timing of human chromosome 6 (62); as well as *XIST* Repeats D, E and F, and its “PfIMI” fragment, both of which are important for *XIST*’s ability to recruit the Polycomb Repressive Complexes, and which associate with HNRNPK and HNRNPU by eCLIP ((19, 20); Figure 3B). Overall, query domains from *XIST* exhibited an enrichment towards hits in presumed repressive lncRNAs, as might be expected given its repressive function (15 *XIST* queries were enriched for hits in repressors, 3 were enriched for hits in activators, and another one was enriched in both classes of lncRNA, out of 21 total queries from *XIST*; Table S9).

As a parallel way to gauge the extent of enrichment in one functional class of lncRNAs over the other, we ranked query domains by their overall difference in hit length between activators and repressors. Across *p* value thresholds for hmSEEKR hit definition, the strongest enrichments among the set of activating lncRNAs included the MLL-associated region within *HOTTIP* (49), the first 1kb of *NEAT1* (22), the proposed P300-recruiting eRNAs produced near the genes *Sp3* and *YY-1* (50), and ncRNA-a7/TRENRA, which has been shown to associate with Mediator ((48); Figure 6); each of these enrichments were highest within the spliced annotations of the set of activating lncRNAs (Figure 6A-C). Hits to interval 5 of the HNRNP-binding domain *ASAR6-141* and several fragments of *NEAT1* were also enriched at all *p* value thresholds within nascent annotations of activating lncRNAs (Figure 6A-C). In contrast, among the queries most enriched in repressive lncRNAs were intervals 2 and 4 of the HNRNP-binding domain of *ASAR6-141* (62), intervals 8 and 11-14 from *XIST*, which lack a clearly ascribed function but by analysis of eCLIP data, robustly interact with the HNRNP or HNRNP-like proteins HNRNPU, SAFB, and KHDRBS1, as well as intervals 4,7, 21, and 22 of *NEAT1*, which interact with a diverse set of proteins by eCLIP (Figures 6A-C and 3B). Across all *p* value thresholds, no single query was enriched in the sets of activating or repressive lncRNAs by more than ∼20% of its aggregate hit length, respectively (Figure 6A-C; Table S9). Among nascent annotations, we observed no significant correlation between percent difference in hit length and GC content of query domains, and among spliced annotations, we observed significant positive correlations (Figure 6D-F).

**Figure 6.**
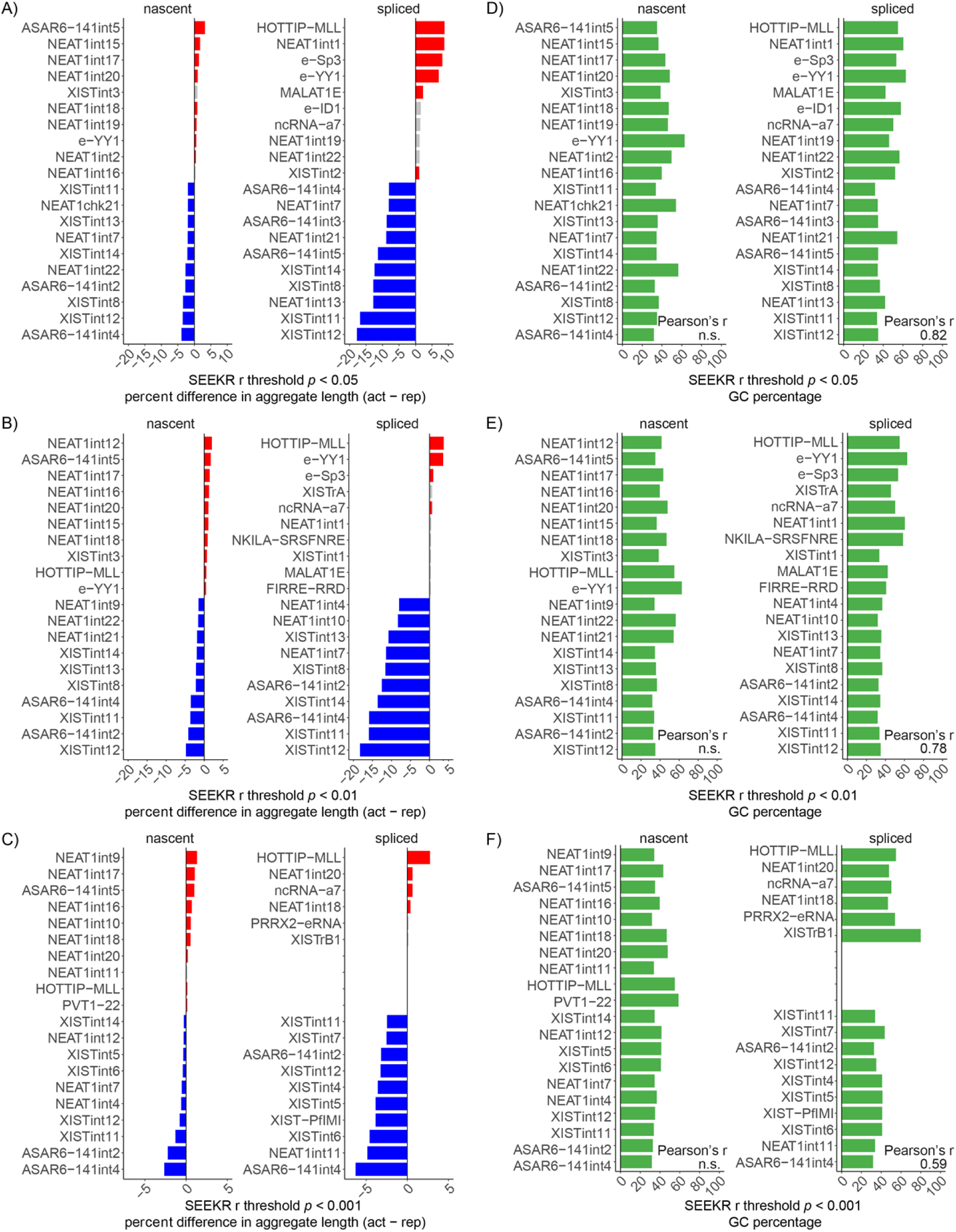
hmSEEKR identifies lncRNAs with local similarities to known regulatory domains. (A, B, **C)** Top 10 query domains whose sum of hit lengths exhibited the greatest percent differences between the sets of presumed activating and repressive lncRNAs, at the *p* < 0.05, *p* < 0.01, and *p* < 0.001 thresholds for hmSEEKR hit definition. Red and blue bars, query domains significantly enriched in the activating or repressive lncRNA sets; grey bars, domains not significantly enriched in either set (Fisher’s exact). **(D, E, F)** GC-percentages of lncRNA query domains listed in (A, B, C). Pearson’s r values examining correlation of GC percentage versus percent difference is shown on the bottom right of each panel. “n.s.”, no significant correlation. All r values displayed represent significant correlations (*p* ≤ 0.01). All searches performed at *k*-mer length *k* = 6. See also Tables S7-9.

Thus, within the set of presumed activating lncRNAs, domain-based searches identified enrichments of lncRNA fragments proposed to interact with several transcriptional effector complexes, and within the set of presumed repressive lncRNAs, domain-based searches identified enrichments of lncRNA fragments shown to bind HNRNP proteins. No single domain exhibited exclusive hits in one set of lncRNAs or the other, supporting the view that the mere presence of a single domain is likely insufficient to ascribe regulatory function to a lncRNA. The domains that were the most enriched within spliced annotations of presumed activating lncRNAs generally had a higher GC content than those that were enriched among presumed repressive lncRNAs.

## Discussion

We present hmSEEKR, a *k*-mer based HMM that enables transcriptome-wide scans for RNA domains that harbor nonlinear similarity to a query domain of interest. The algorithm builds on the SEEKR framework to enable detection of regional similarities without predefined knowledge of where similarities might be located within the RNAs being searched (6–8). To detect similarity, hmSEEKR requires that the query domain(s) differ in *k*-mer content from the set of sequences used as background. In our uses of hmSEEKR thus far, this requirement has been easily met by selecting large sets of sequences as background, such as the entire lnc-ome of the organism in question (16). hmSEEKR also requires that end-users provide parameters that define the likelihood of transitioning between the query and null states, which can be determined through empirical evaluation of hits across a grid of transition parameters. In our studies above, we performed hmSEEKR grid searches using 78 different query domains (Table S1). From those searches, we selected as defaults the two transition parameters that we found to be the most generalizable, enabling end-users to perform preliminary hmSEEKR scans without the need to perform computationally intensive grid searches. However, to perform rigorous custom scans, we recommend that end-users carry out grid searches using their own custom query domains and background sequences. If computational resources are limiting, we would suggest training with randomly selected subsets of lncRNA sequences rather than entire transcriptomes; the larger the randomly selected subset, the more stable the output data associated with each set of tested transition parameters, but the greater the computational cost.

As evidence that hmSEEKR can detect meaningful similarities within transcriptomes, we demonstrated that the algorithm can successfully identify randomly inserted query fragments (Figure 2), it can identify hits that exhibit above-background levels of association with similar sets of proteins as their corresponding query domains (Figure 3), and, in sequential and combinatorial domain scans, it can identify RNAs whose overall networks of protein association are more similar to those of their corresponding query lncRNA than would be expected by chance (Figure 5).

In applications of hmSEEKR, at our highest significance thresholds for hmSEEKR hit definition (*p* < 0.001), we observed that many domains within *XIST*, *NEAT1*, and *MALAT1* were rare within the K562-expressed, chromatin-associated transcriptome, exhibiting a significant hit to less than 200 other transcripts or to their own transcript only. These observations support the view that *XIST*, *NEAT1*, and *MALAT1* are generally distinct from other expressed transcripts, consistent with their known distinct subnuclear localization and biological roles.

However, at lower significance thresholds for hmSEEKR hit definition (*p* < 0.05), we found that many domains within the lncRNAs *XIST*, *NEAT1*, and *MALAT1* exhibit prevalent similarity to transcripts produced from both protein-coding and lncRNA genes, frequently within intronic regions. Relatedly, combinatorial scans for sequential similarity to minimal versions of *XIST* and *NEAT1* almost exclusively identified intron-containing/nascent transcript isoforms as being the most similar. These results echo those from our recent analyses of RNA-protein associations in mouse trophoblast stem cells, which showed that protein association patterns in the repressive lncRNAs *Xist*, *Airn*, and *Kcnq1ot1* exhibited similarities to transcripts produced from both lncRNA and protein-coding genes, and particularly for *Airn* and *Kcnq1ot1*, that the majority of similar transcripts were intron-including or nascent isoforms (14). While the functional relevance of these similarities remains to be determined, our data support the view that intronic elements represent a possible reservoir of regulatory domains that have functions beyond splicing control. Intronic sequences may be most relevant to lncRNAs, which as a class, are known to be spliced inefficiently (106–109). However, prior to export chromatin, even RNAs produced from protein coding genes may exert lncRNA-like regulatory functions, either as nascent transcripts or as spliced products if RNA export after splicing is delayed (119–128). A recent study found that the *TTN* pre-mRNA can form an architectural hub that regulates gene expression, splicing, and chromatin, an example of regulatory RNA produced from a protein-coding gene (129). In the future, it will be interesting to examine the potential regulatory functions of *XIST*- and *NEAT1*-similar transcripts, the majority of which detected in this study were produced from protein-coding genes. In that regard, our catalogue of expression correlations across GTEx samples may prove useful (47); we identified several *XIST*- and *NEAT1*-similar transcripts that exhibit significant positive and negative expression correlations with nearby genes, consistent with the possibility of RNA-mediated cis-regulatory effects.

In a separate application of hmSEEKR, we examined the extents to which previously identified RNA regulatory domains exhibited regional enrichments within lncRNAs linked to the local activation and repression of transcription in cis. We identified an MLL-associated domain of *HOTTIP*, and a promoter-proximal domain of *NEAT1* and two enhancer RNAs (*e-Sp3* and *e-YY1*), as the most significantly enriched domains within the set of activating lncRNAs (22, 49, 50). In contrast, HNRNP-binding fragments from *XIST* and the lncRNA *ASAR6* were the most significantly enriched within the set of repressive lncRNAs (62). The enrichments of these domains were generally modest, exhibiting on the order of a few percentage points difference in aggregate hit lengths between activating and repressive lncRNAs. The modest enrichments could be viewed as support for the idea that many lncRNAs may carry out gene regulatory roles through the combined action of multiple functional domains. Other explanations may also contribute to the modest enrichments, including the likelihood that hmSEEKR gives rise to false positive and false negative detections, and the likelihood that both the functional domains used as search features and the sets of lncRNAs being searched contain false positives. Disentangling the complex relationships between lncRNA sequence and function remains a major challenge, and our domain-centric analyses provide sequence-based footholds that can be used to guide future investigations.

By extension, *k*-mer-based comparisons tend to be lower in stringency than more traditional forms of linear sequence alignments. For that reason, we view hmSEEKR primarily as a hypothesis generating tool. The mere presence of regional *k*-mer similarity is, by itself, insufficient to confidently ascribe function to any RNA. However, knowledge of where regional *k*-mer similarities might exist can enable functional interrogation in ways that are informed by underlying sequence information. Of equal importance, hmSEEKR can be trained on a single query domain and in most cases, can be effectively used on a personal computer. For these reasons, hmSEEKR represents an important complement to emerging machine learning models, which, particularly where lncRNAs are concerned, require training corpora that do not yet exist and high-performance computing resources that are not broadly accessible. Beyond lncRNAs, we envision that hmSEEKR will also be useful for identifying important features within DNA regulatory elements and the untranslated regions of mRNAs, both of which can encode functions through degenerate sequences. hmSEEKR is available for download through the Python Package Index, the Docker Hub, and GitHub (https://github.com/CalabreseLab/hmseekr). A separate GitHub page contains the code used to generate the figures in this manuscript as well as additional example uses of hmSEEKR (https://github.com/CalabreseLab/hmseekr_manuscript).

## Acknowledgements

This work was supported by NSF grant DBI-2228805; NIH grants R01GM136819, R01GM121806, and R35GM153293; and the Yang Family Biomedical Scholar Fund (J.M.C.). D.A.S. was supported by T32GM007040. Q.E.E. was supported by F31HG014413. S.B.P. was supported by T32GM135128. A.L. was supported by R01HL111527 and R35GM140844. The sponsors/funders played no role in the study design, data collection and analysis, decision to publish, or preparation of the manuscript. NSF DBI: https://www.nsf.gov/bio/dbi/about, National Institute of General Medical Sciences (NIGMS): https://www.nigms.nih.gov, National Human Genome Research Institute (NHGRI): https://www.genome.gov, UNC RNA Discovery Center:https://unclineberger.org/rnadiscoverycenter/

## Author Contributions

S.L.: Conceptualization, Data Curation, Formal Analysis, Investigation, Methodology, Software, Visualization, Writing – Original Draft Preparation, Review & Editing. D.A.S.: Conceptualization, Methodology, Software, Writing – Review & Editing. Q.E.E.: Data Curation, Formal Analysis, Investigation, Methodology, Software, Writing – Review & Editing. S.P.B.: Methodology. A.L. Conceptualization, Writing – Review & Editing. J.M.C.: Conceptualization, Data Curation, Formal Analysis, Funding Acquisition, Investigation, Methodology, Project Administration, Supervision, Validation, Visualization, Writing – Original Draft Preparation, Writing – Review & Editing.

## Methods

### hmSEEKR

The hmSEEKR package can be found at https://github.com/CalabreseLab/hmseekr, and code used to generate all figures and tables at https://github.com/CalabreseLab/hmseekr_manuscript.

hmSEEKR can be installed via the Python Package Index, via GitHub, or via the Docker Hub (instructions found https://github.com/CalabreseLab/hmseekr). The set of sequences used as the null model in all hmSEEKR runs and grid searches were the set of lncRNA transcripts downloaded from the GENCODE v47 collection and filtered to include only de-duplicated isoforms with “canonical” labels that were greater than or equal to 500nt in length (7). Transition parameters selected for all hmSEEKR scans are listed in Table S1.

The following describes our heuristic to identify transition parameters. Examining outputs of hmSEEKR scans using query domains within *XIST*, *NEAT1*, and *MALAT1*, we observed that transition probabilities ranging from 0.9 to 0.99 yielded hit sequences of length and sequence content that most closely resembled their corresponding queries. Thus, all grid searches performed for this work tested all combinations of query and null transition probabilities ranging from 0.9 to 0.99 with a step of 0.03. For queries whose length was less than 100nt, we set the minimal hit length as 25nt and maximal hit length as 800nt. For queries whose lengths ranged between 100nt and 200nt, we set the minimal hit length as 50nt and maximal hit length as 1000nt. For queries whose length rangedbetween 200nt and 800nt, we set the minimal hit length as approximately one fourth the query length and maximal hit length as approximately 4 times the query length. For queries with length greater than 800nt, we set the minimal acceptable hit length as 200nt and maximal acceptable hit length as 6000nt. For the HSat3_sense and HSat3_anitsense sequences, we set minimal hit length as 25 and maximal as 6000 to better capture the signature of short repeats. In general, we selected qT and nT pairs that yielded the highest median SEEKR r value from the top 50 hit sequences ranked by SEEKR r value, and that yielded a total number of qualified hits after length filtering that were greater than the smaller of 10000 or the median hit number across all qT and nT combinations. Grid searches for all queries were performed at *k*-mer lengths *k* = 4, 5, and 6.

### Defining a set of K562-expressed, chromatin-associated RNAs

To identify a set of chromatin-associated RNAs expressed in K562 cells, we downloaded the GENCODE v47 comprehensive GTF and to it added a single intron-containing isoform for every gene that spanned from its earliest start and ranged to its latest end. The one exception was for the lncRNA *KCNQ1OT1*, whose single transcript we adjusted to be exclusively mono-exonic and spanning hg38 coordinates chr11:2,608,328-2,699,994 on the negative strand, consistent with our prior investigations of RNA-seq read density in K562 cells (7). Using this transcript collection, we then used kallisto to align total RNA-seq (ENCSR885DVH), fractionated RNA-Seq (chromatin: ENCSR000CPY; PolyA cytosolic: ENCSR000COK), and Bru-Seq (ENCSR729WFH), and Bru-Chase (2hr: ENCSR633UIR; 6hr: ENCSR762OPQ) data collected in K562 cells (1, 25). The transcripts whose average TPM values from total RNA-seq were greater than 0.125, whose [chromatin_TPM]/([chromatin_TPM]+[cytosolic_TPM]) values were greater than 0.75, and whose lengths were greater than or equal to 500nt were retained as expressed and chromatin-associated. To further reduce redundancy with our set of transcripts, for each gene producing a chromatin-associated RNA, we retained at maximum, only a single intron-excluding (i.e. “spliced”) isoform and a single intron-including (i.e. “nascent”) isoform. Some genes produced only intron-excluding isoforms detectable as chromatin-associated, others produced only intron-including isoforms detectable as chromatin-associated, and others produced both intron-excluding and intron-including isoforms. In total, we retained 8630 intron-excluding isoforms covering ∼17 million nucleotides of sequence, and 6663 intron-including transcript isoforms covering ∼253 million nucleotides of sequence. Bru-Chase-seq estimates of stability in Figure 5 were calculated as [Bru-Chase-6hr]/([Bru-Chase-0hr]+0.125).

### Recovery of randomly inserted query domains from the chromatin-associated transcriptome

To perform the analyses displayed in Figures 2B and 2C, we inserted each *XIST*, *NEAT1*, and *MALAT1* query domain randomly among the set of K562-expressed, chromatin-associated transcripts 1000 times, then used hmSEEKR and the python code outlined in https://github.com/CalabreseLab/hmseekr_manuscript to measure the recovery of the inserted queries at *k*-mer lengths *k* = 4, 5, and 6. In Figure 2A and B, within each mutational frequency bin, ranging from 0% (WT) to 50%, the data from all *XIST*, *NEAT1*, and *MALAT1* query domains are plotted together. Mutations were made by randomly selecting nucleotides within each query and changing it to one of the other three nucleotides, as documented in https://github.com/CalabreseLab/hmseekr_manuscript.

### eCLIP data processing

All per-replicate eCLIP and seCLIP BAM files were obtained from ENCODE for the K562 cell line, that were GRCh38-aligned, filtered alignments with a ‘released’ status (24). For RBP targets with both seCLIP (single-end, forward stranded) and eCLIP (paired-end, reverse stranded) experiments, the seCLIP experiment was used (36, 130). The control experiments corresponding to each targeted eCLIP experiment were also downloaded. To identify the region most proximal to the site of the bound RBP, the first read of each aligned pair in all paired-end experiments was discarded (second read retained by samtools view -f 131) which also enabled all alignments to be processed as if single-ended and forward stranded for the remainder of the analysis (131). Alignments of replicated experiments were downsampled with respect to the replicate with the fewest aligning reads as identified through samtools view -c, and reads were retained with samtools view -b- s <downsampling-fraction> where the downsampling fraction was [1 – (replicate 1 read counts – replicate 2 read counts)/(minimum read count between replicates)] (7, 131).

Downsampled replicates were merged and the final number of reads were calculated with samtools view -c. Merged alignments were filtered for quality and split by strand so that samtools view -b -q 30 -f 144 extracted reads originating from genes on the negative strand for eCLIP experiments and samtools view -b -q 30 -f 160 extracted reads originating from genes on the positive strand. For seCLIP experiments, samtools view -b -q 30 -f 16 extracted the negative stranded reads and samtools view -b -q 30 -F 16 extracted the positive stranded reads. The same process was followed to split control files by strand. For each experiment, called peaks were downloaded from ENCODE and reads aligning to the peak regions were retained in both the experimental replicates and their respective paired control alignments with bedtools intersect -s - split -ubam (132). For experiments with replicates, reads aligned to peak regions for either replicate were retained in the final alignment. In the region-by-region analyses of eCLIP data in Figure 3, each eCLIP protein target was paired with its respective control dataset. In the network analyses of Figure 5, a single control dataset was generated by merging all reads under peaks from each eCLIP-paired control dataset used; this single control dataset was used in all background-correction operations.

### eCLIP enrichment under XIST, NEAT1, and MALAT1 query domains and hmSEEKR hits

The multicov function from BEDtools and the merged bams for each eCLIP protein target along with the merged control described in the “eCLIP data processing” section above were used to count eCLIP reads under each *XIST, NEAT1,* and *MALAT1* query domain and all hmSEEKR hits (132). We retained protein targets whose reads-per-million (RPM)-normalized eCLIP density was at least two-fold greater than the RPM-normalized density of control and whose input-control normalized read densities were > 2. Among the protien targets that passed these thresholds, the top five protein targets for each query domain (ranked by eCLIP/control ratio) are shown in Figure 3B. In *MALAT1* query fragment #9, only a single protein target, SF3B4, had an eCLIP/control ratio of greater than two.

To generate the *p* values indicated by red shading in Figure 3B, we compared the list of control-corrected eCLIP signal values under each hmSEEKR hit (defined as [eCLIP-RPM minus control-RPM]) to a paired list of control-corrected eCLIP signal values that were calculated by using BEDtools to shuffle the position of each hmSEEKR hit once within the set of coordinates of genes producing K562-expressed, chromatin-associated RNAs, retaining the length of the original hit (132). We then used Wilcoxon paired test and Benjamini-Hochberg correction to compare whether hmSEEKR hits had significantly higher control-corrected eCLIP signal values than their shuffled and paired controls.

To assess significance of combinatorial enrichment under hmSEEKR hits derived from the queries listed in Figure 3C, we first created two separate lists of shuffled controls for each query, defined as the satellite and anchor controls. For each set of protein targets examined, we counted the number of times in which the input-control normalized read densities for hmSEEKR hits were greater than the input-control normalized read densities in the paired anchor controls, for both (or all three) proteins being analyzed. Separately, we counted the number of instances in which input-control normalized read densities for the satellite controls were greater than the input-control normalized read densities in the paired anchor controls, again, for both (or all three) proteins being analyzed. We then used Fisher’s exact test in a 2x2 contingency table to determine if hmSEEKR hit regions had significantly more instances in which all of the proteins being analyzed under the query had signal greater than their corresponding pairs of satellite and anchor controls.

### Intersection of hmSEEKR hits with gene types and gene features

Gene types and features were defined using all genes and transcripts in the GTF from the GENCODE v47 comprehensive primary assembly of GRCh38. The coordinates of all genes, exons, UTRs (5’ and 3’), and CDS regions were used as documented in the GTF. Additional features were annotated, including transcriptional start and end sites (TSS, TES), 5’ and 3’ splice sites, and proximal and distal introns. The region within 1,000 nucleotides of the 5’ end or 3’ end of a gene were categorized as a TSS or TES, respectively. Splice sites were defined as the regions within 50 nucleotides of junctions between exons and introns, with the 5’ splice site following an upstream exon and the 3’ splice site preceding a downstream exon. Regions of introns that were greater than 2,000 nucleotides from an exon were classified as distal introns, and intronic regions closer to exons were defined as proximal introns. For any spanning feature near the end of a chromosome, the bounds of the chromosomes truncated the region.

To calculate significance of hit enrichment within gene types and gene features in Figures 4A and 4B, the list of hmSEEKR hits from each query was shuffled three times within the coordinates of the set of genes producing K562-expressed, chromatin associated RNAs using BEDtools (132). The overlap between each feature and each hmSEEKR hit and shuffled hit was calculated using BEDtools. The average of the feature overlap value across the three sets of shuffled coordinates was retained. Fisher’s exact test was used to determine if the hmSEEKR hits were associated with the feature to a greater degree than the shuffled controls. The Benjamini-Hochberg correction was applied to each genetype/genefeature across all queries to obtain adjusted *p* values.

### Sequential similarity searches for matches to minimal XIST and NEAT1

To identify K562-expressed, chromatin-associated RNAs with sequential similarity to the sets of minimal functional domains within *XIST* and *NEAT1*, we firstly ran individual or combinatorial (seqs-to-summary) searches on the following domains: *XIST* Repeat A, D, E, F, and B1 and B2 (seqs-to-summary); and *NEAT1* interval N1, N9 and N10 (seqs-to-summary), and N11 through N16 (seqs-to-summary). For each host lncRNA searched (*XIST* and *NEAT1*), we retained only those K562-expressed, chromatin-associated RNAs that had an hmSEEKR hit to all of the functional modules searched. Then we employed the following sequential logic. For transcripts to be considered *XIST*-similar, we required that they have hits to Repeat A and Repeat F within their first 3kb, followed by hits to Repeat B1/B2 and Repeat D (in either order), followed by a hit to Repeat E. For transcripts to be considered *NEAT1*-similar, we required that they have a hits to N1 within its first 3kb, followed by hits to N9-10, followed by at least 1kb of hits to N11-16, and lastly, a hit to N22.

### Comparing protein association networks between transcripts

To establish protein association networks to compare between *XIST* and *NEAT1* and their respective groups of sequentially similar transcripts, we first collated a list of proteins that across all K562 eCLIP datasets, were within the top five most-enriched over each of the relevant query domains in *XIST* and *NEAT1*, and whose hmSEEKR hits in individual searches exhibited associations that were significantly greater than randomly shuffled controls across the K562-expressed, chromatin-associated transcriptome (shaded boxes in Figure 3B). For this set of proteins (21 in the *XIST* set, and 17 in the *NEAT1* set), we retained eCLIP read density and control read density under peaks using an approach analogous to that described in (14). Specifically, eCLIP reads under peaks were identified as described above in the section “*eCLIP data processing*”. BEDtools multicov was then used to calculate replicate-averaged, RPM-normalized read densities in 25nt bins over every gene producing a K562-expressed, chromatin-associated RNA (132). Then, in each bin, RPM signal-under-peak for each eCLIP target minus corresponding control was calculated, and any negative values were set to zero.

Once binned and background-corrected values were obtained, networks of protein association across each gene producing a K562-expressed, chromatin-associated RNA were calculated in the following manner. For each gene, we calculated Pearson’s r values comparing control-corrected read densities in 25nt bins for all possible pairwise comparisons of each protein in each list of *XIST*- and *NEAT1*-associated proteins. In cases where all control-corrected read densities for a given protein i were comprised of zero values, Pearson’s comparisons involving that protein were set to a value of ’0’. Genes whose control-corrected read densities were comprised of zero values across all proteins in a list were removed. The list of Pearson’s r values between all possible pairwise comparisons of control-corrected eCLIP densities within a gene is referred to as its protein association network. After calculating within-gene protein association networks, we then performed between-gene network comparisons, again using Pearson’s correlation. Wilcoxon rank-sum test was used to calculate the significance of the difference between the protein association networks of *XIST*- and *NEAT1*-similar genes and the networks from all genes producing K562-expressed, chromatin-associated RNAs (Figure 5C).

### Analysis of GTEx data

GTEx data were dowloaded from https://gtexportal.org/home/downloads/adult-gtex/bulk_tissue_expression (GTEx_Analysis_2026-05-19_v11_RNASeQCv2.4.3_gene_reads.gct) and metadata were downloaded from https://gtexportal.org/home/downloads/adult-gtex/metadata

(GTEx_Analysis_v11_Annotations_SampleAttributesDS.txt).

Selected tissue types in the SMTSD columns were included for DESeq2 analysis for differentially expressed genes. The selected tissue types were: “Cells - Cultured fibroblasts”, “Whole Blood”, “Spleen”, “Cells - EBV-transformed lymphocytes”, “Lung”, “Liver”. In total, these tissues included 2925 samples. Genes were excluded from DESeq2 analyses if they did not have at least 1 count in at least 1500 samples. Cells - Cultured fibroblasts was used as the reference level for the DESeq2 model. Differential expressed genes (DEGs) were defined as having adjusted *p* values of < 0.05 and log2-fold changes of > 1. DEGs from all tissue types compared to fibroblasts were merged together to form a final DEGs list. The correlations between the expression of nearby genes and *XIST*-similar genes (*p* < 0.05) and *NEAT1*-similar genes (*p* < 0.01) were further analyzed if the *XIST-* and *NEAT1*-similar were in the final DEGs list. For each retained *XIST*- and *NEAT1*-similar DEG, its Pearson’s correlation with all other DEGs on the same chromosome and within +/- 500kb of its transcription start site were calculated using DESeq Normalized counts. Gene pairs that had Pearson’s correlation >0.25 or <-0.25 and adjusted *p* values of < 0.05 were saved and reported in Table S6.

### Domain analyses in activating and repressive lncRNAs

hmSEEKR searches were performed using the domains listed in Tables S6 and S8 to search the intron-containing/nascent and spliced annotations within the sets of presumed activating and repressive lncRNAs listed in Table S8. The summed length of hits for each query domain in each set of lncRNAs searched was calculated at each *p* value threshold for hmSEEKR hit definition (*p* < 0.05, 0.01. and 0.001). This sum and the sum of length of non-matching sequence, each divided by 100, was used to populate a 2x2 contingency table and perform subsequent Fisher’s exact tests.

---

**Figure S1.**
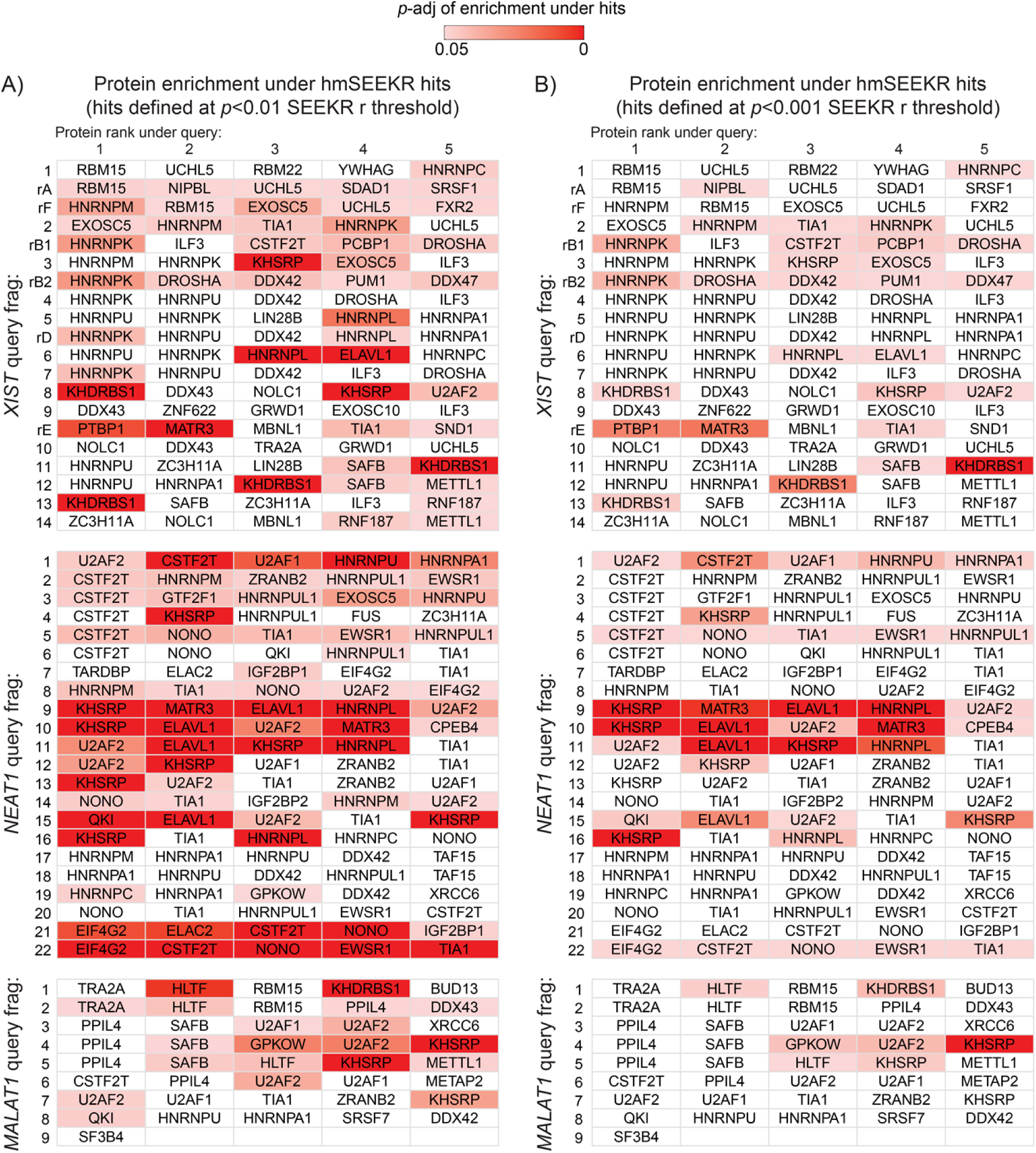
Protein enrichment under hits to query domains at *k*-mer length *k* = 6 and thresholds for hmSEEKR hit definition of *p* < 0.01 (A) and < 0.001 (B). Each row in (A) and (B) corresponds to a query domain and is displaying the Wilcoxon signed-rank test, Benjamini-Hochberg adjusted *p* value of CLIP enrichment for each of that query’s top five most enriched proteins underneath the hits to the query. Related to Figure 3.

**Figure S2.**
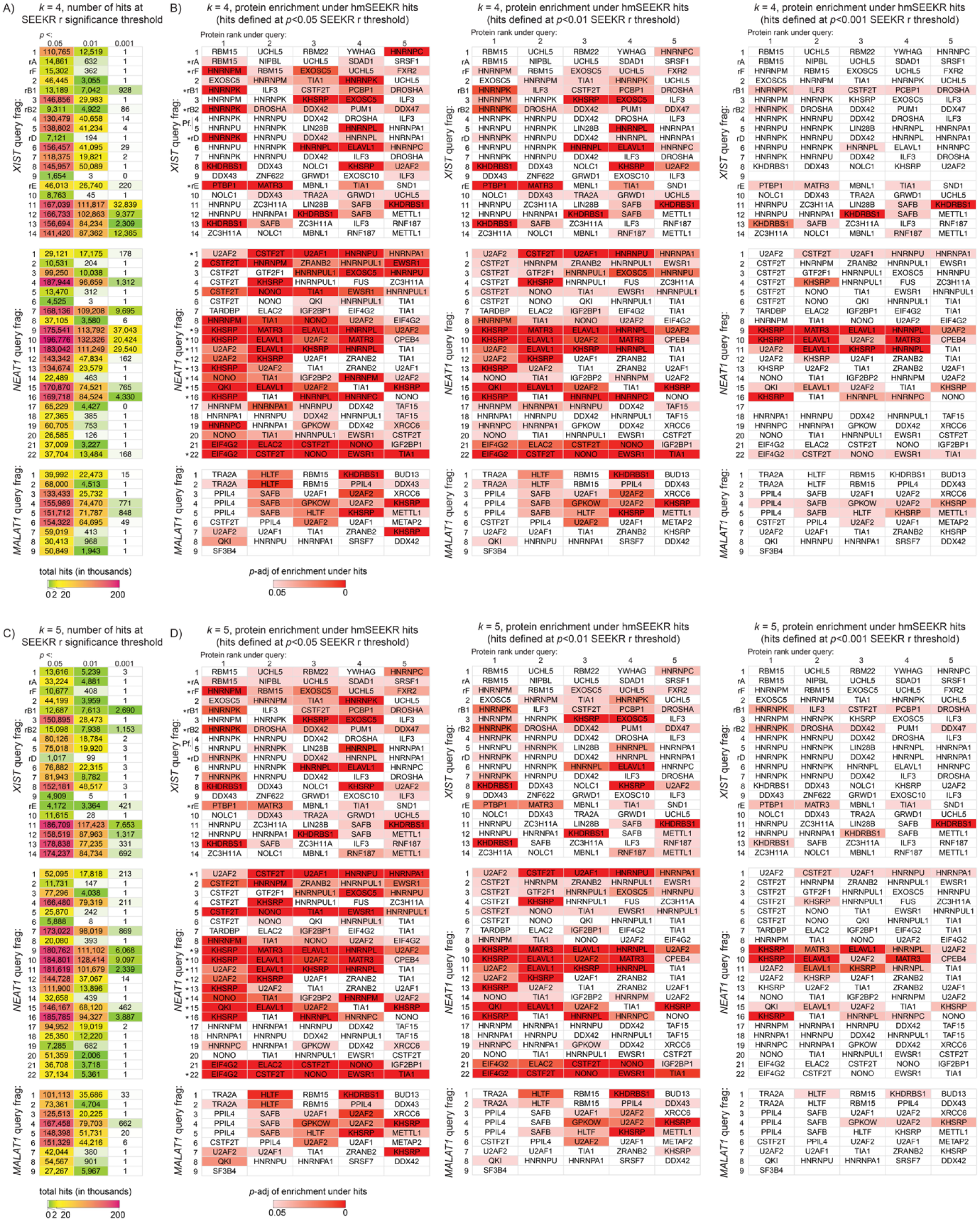
Location of hmSEEKR hits and enrichment of CLIP data under hits with searches performed at *k*-mer length *k* = 4 and 5. **(A)** At *k*-mer length *k* = 4, number of hits to each *XNM* query in the set of genes that produce chromatin-associated RNAs in K562 cells. **(B)** At *k*-mer length *k* = 4, significance of enrichment of CLIP signal under hmSEEKR hits versus shuffled controls for query-associated RBPs across all *XNM* query domains (Wilcoxon signed-rank test, Benjamini-Hochberg adjusted), at the *p* < 0.05, 0.01, and 0.001 thresholds for hit definition. **(C)** and **(D)**, same as (A) and (B), but for searches performed at *k*-mer length *k* = 5. Related to Figure 3.

**Figure S3.**
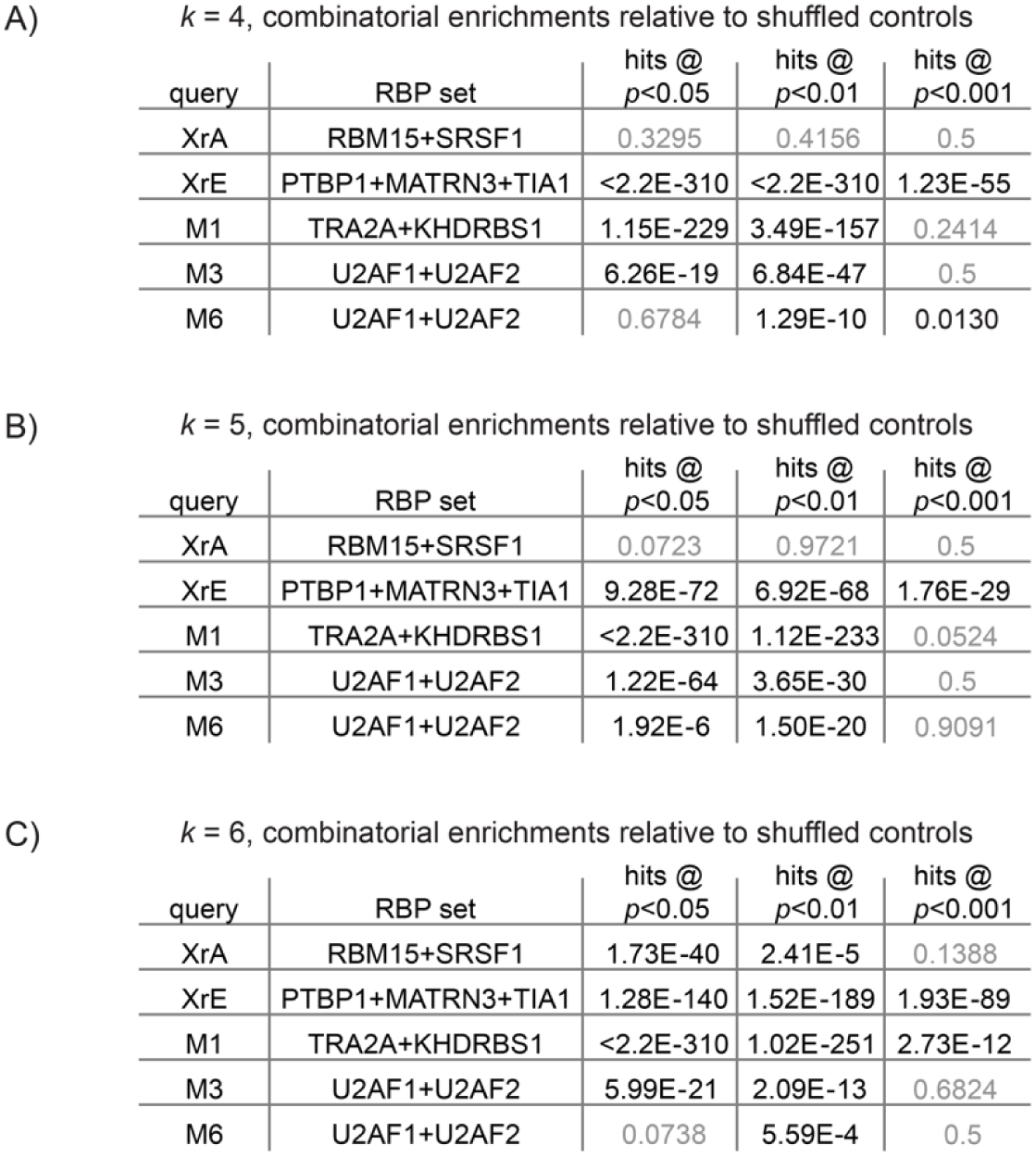
Significance of combinatorial enrichments of CLIP signal under hmSEEKR hits for *XIST* and *MALAT1* query domains that have been previously shown to interact with more than one protein client. Significance of combinatorial enrichment (Fisher’s exact) is shown at each of three separate *p* value thresholds for hmSEEKR hit definition. Data shown for searches performed using *k*-mer lengths *k* = 4 **(A)**, 5 **(B)**, and 6 **(C)**. Related to Figure 3.

**Figure S4.**
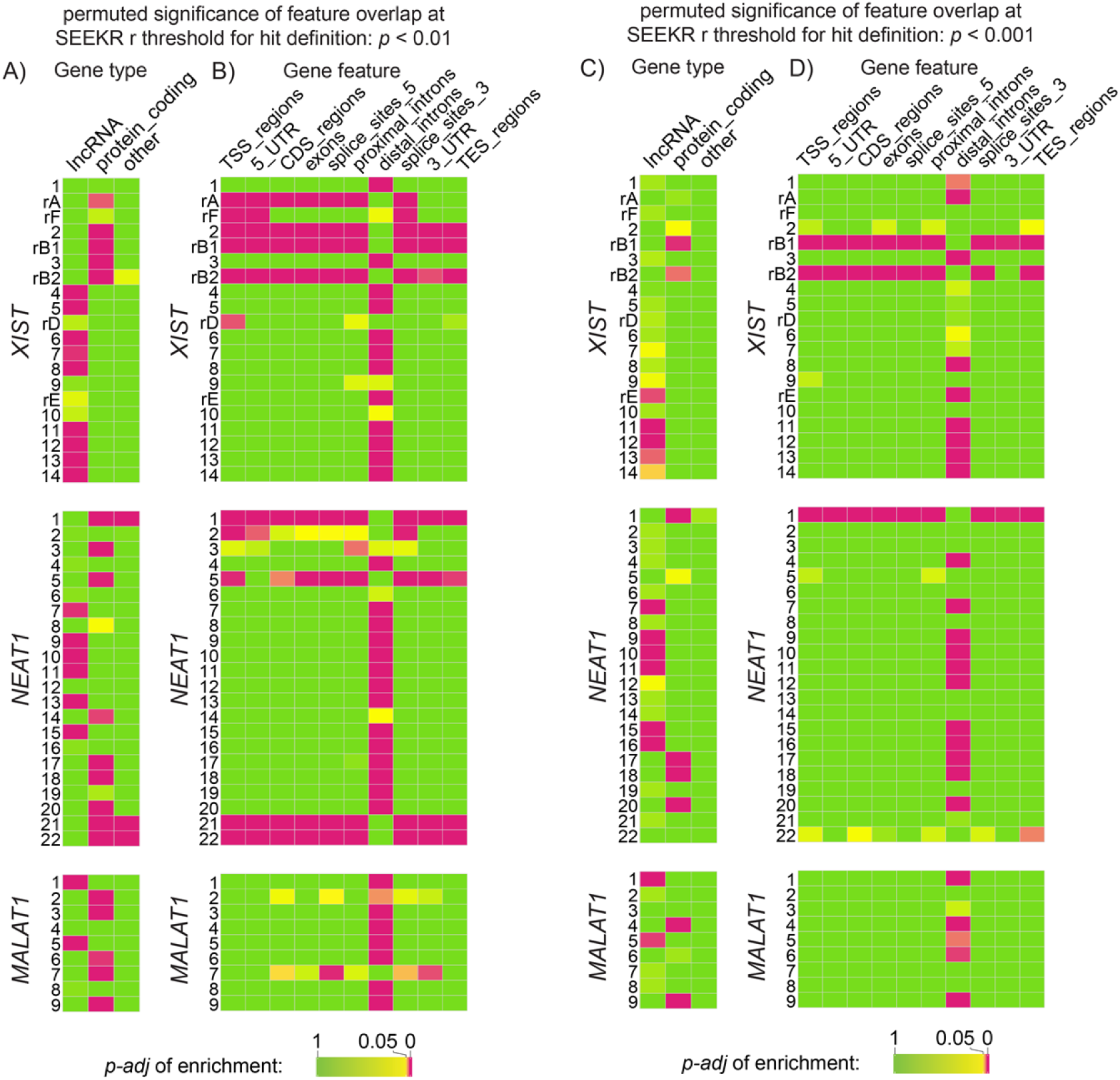
Gene-centric overlaps of hmSEEKR hits at *k*-mer length *k* = 6 and thresholds for hmSEEKR hit definition of *p* < 0.01 and *p* < 0.001. (**A**) For each *XNM* query, adjusted *p* value of hit enrichment in the set of genes that produce chromatin-associated RNAs in K562 cells (Fisher’s exact, Benjamini-Hochberg adjusted); and (**B**) *p* value of hit enrichment in the features of genes that produce chromatin-associated RNAs in K562 cells (Fisher’s exact, Benjamini-Hochberg adjusted). Hits were defined using a SEEKR-derived *p* value threshold of *p* < 0.01. (C) and (D), same as (A) and (B) but using a SEEKR-derived *p* value threshold of *p* < 0.001. Related to Figure 4.

